# HDAC and MAPK/ERK Inhibitors Cooperate to Reduce Viability and Stemness in Medulloblastoma

**DOI:** 10.1101/521393

**Authors:** Mariane da Cunha Jaeger, Eduarda Chiesa Ghisleni, Paula Schoproni Cardoso, Marialva Siniglaglia, Tiago Falcon, André T. Brunetto, Algemir L. Brunetto, Caroline Brunetto de Farias, Michael D. Taylor, Carolina Nör, Vijay Ramaswamy, Rafael Roesler

**Affiliations:** Cancer and Neurobiology Laboratory, Experimental Research Center, Clinical Hospital (CPE-HCPA), Federal University of Rio Grande do Sul, Porto Alegre, RS, Brazil; Children’s Cancer Institute, Porto Alegre, RS, Brazil; Bioinformatics Core, Experimental Research Center, Clinical Hospital (CPE-HCPA), Federal University of Rio Grande do Sul, Porto Alegre, RS, Brazil; The Arthur and Sonia Labatt Brain Tumour Research Centre, The Hospital for Sick Children, Toronto, ON, Canada; Developmental and Stem Cell Biology Program, The Hospital for Sick Children, Toronto, ON, Canada; Department of Laboratory Medicine and Pathobiology, University of Toronto, Toronto, ON, Canada; Division of Neurosurgery, The Hospital for Sick Children, Toronto, ON, Canada; Division of Haematology/Oncology, The Hospital for Sick Children, Toronto, ON, Canada; Department of Pharmacology, Institute for Basic Health Sciences, Federal University of Rio Grande do Sul, Porto Alegre, RS, Brazil

**Keywords:** Histone deacetylase, Extracellular-regulated kinase, Stem cells, Medulloblastoma, Brain tumor

## Abstract

Medulloblastoma (MB), which originates from embryonic neural stem cells (NSCs) or neural precursors in the developing cerebellum, is the most common malignant brain tumor of childhood. Recurrent and metastatic disease is the principal cause of death and may be related to resistance within cancer stem cells (CSCs). Chromatin state is involved in maintaining signaling pathways related to stemness, and inhibition of histone deacetylase enzymes (HDAC) has emerged as an experimental therapeutic strategy to target this cell population. Here, we observed antitumor actions and changes in stemness induced by HDAC inhibition in MB. Analyses of tumor samples from patients with MB showed that the stemness markers *BMI1* and *CD133* are expressed in all molecular subgroups of MB. The HDAC inhibitor (HDACi) NaB reduced cell viability and expression of *BMI1* and *CD133* and increased acetylation in human MB cells. Enrichment analysis of genes associated with *CD133* or *BMI1* expression showed mitogen-activated protein kinase (MAPK)/ERK signaling as the most enriched processes in MB tumors. MAPK/ERK inhibition reduced expression of the stemness markers, hindered MB neurosphere formation, and its antiproliferative effect was enhanced by combination with NaB. These results suggest that combining HDAC and MAPK/ERK inhibitors may be a novel and more effective approach in reducing MB proliferation when compared to single-drug treatments, through modulation of the stemness phenotype of MB cells.

## Introduction

Medulloblastoma (MB), the most common malignant pediatric brain tumor, likely arises from genetic and epigenetics abnormalities during cerebellar development (Wang and Wechsler-Reya 2014; Vladoiu et al. 2019). Integrative genomic, epigenomic and transcriptional analyses have shown that MB is not a single tumor type. Instead, it is comprised of at least four molecular subgroups: Wingless (WNT), Sonic hedgehog (SHH), Group 3 and Group 4, that also have distinct demographic, clinical, and prognostic features (Taylor et al. 2012). The WHO 2016 classification of central nervous system (CNS) tumors divides MB into WNT-MB, SHH-MB/*TP53* wild type, SHH-MB/*TP53* mutated, Group 3 and Group 4 (Louis et al. 2016). This molecular identification of MB subgroups has resulted in several clinical trials of subgroup specific therapies, however, except for de-escalation of therapy for WNT patients, there are a paucity of new treatment options for most patients. Specifically, the long-term neurocognitive sequelae of therapy affect the life quality of MB survivors while patients with recurrent and metastatic disease usually succumb to their disease (Schwalbe et al. 2017; Northcott et al. 2019).

According to the cancer stem cell (CSC) hypothesis, recurrence and metastasis in some tumor types may arise from a specific tumor cell subpopulation, the CSC, a distinct population of cells presenting stem cell-like properties such as clonal long-term repopulation and self-renewal capacity (Kreso and Dick 2014; Peitzsch et al. 2017). In some cancers, including CNS tumors, CSCs drive tumor initiation and progression and are able to recapitulate the phenotype of the tumor from which they were derived (Singh et al. 2004). In MB, CSCs are identified by the expression of neural stem cell (NSC) markers such as *CD133, BMI1, Nestin* and *Sox2,* by their capacity to form neurospheres *in vitro,* and by tumorigenicity in animal xenograft models (Hemmati et al. 2003; Singh et al. 2004; Huang et al. 2019). These highly tumorigenic cells display features including stemness, therapeutic resistance, and invasion (Annabi et al. 2008; Hambardzumyan et al. 2008; Lu et al. 2009; Yu et al. 2010; Liu et al. 2015).

Given that a failure in maturation of cerebellar progenitor cells is likely at the origin of MB, and the resistance characteristics presented by CSCs may be responsible for therapeutic failure, modulation of cellular differentiation can be a relevant strategy. In this context, epigenetic modulators have emerged as an exciting therapeutic avenue, particularly histone deacetylase inhibitors (HDACis) (Ververis et al. 2013; Klonou et al. 2018). HDACs are enzymes that control, in a balance with histone acetyltransferases (HATs), histone acetylation regulating chromatin state. HDACis promote the acetylation of histones and non-histone proteins, modify gene expression profiles, and regulate many biological processes involved in tumor homeostasis, such as apoptosis, cell-cycle arrest, angiogenesis, and differentiation (Ververis et al. 2013). HDACi play a role as differentiation inducers in several childhood cancers including neuroblastoma (Frumm et al. 2013), Ewing sarcoma (Souza et al. 2018) and MB (Li et al. 2004; Pak et al. 2019). Expression of HDAC2 is higher in patients with MB with poor prognosis (SHH, Group 3 and Group 4), and *MYC*-amplified MB cell lines show increased mRNA levels of class I HDACs compared to normal cerebellar tissue (Ecker et al. 2015). The antiproliferative effect of HDACi in MB is accompanied by morphological changes, cell cycle arrest, increased expression of glial and neuronal markers, and reduced tumorigenicity (Sonnemann et al. 2006; Spiller et al. 2008; Marino et al. 2011; Nör et al. 2013; Phi et al. 2017; Yuan et al. 2017). We have previously shown that sodium butyrate (NaB), a class I and IIa HDACi, decreases neurosphere formation in MB cultures (Nör et al. 2013). Using different HDACis, a recent study found a similar effect on neurosphere survival, even in the absence of significant changes in the expression of the stemness marker *CD133* (Yuan et al. 2017).

Taking into account that the stemness phenotype is proposed as a poor prognostic marker in many cancer types including MB (Glinsky et al. 2005; Shats et al. 2011), and the effects of HDACi in regulating stemness are still not fully understood, the aim of this study was to investigate how NaB influences aspects of the stemness phenotype of MB cells. We examined the expression of *BMI-1,* a potent inducer of NSC self-renewal and neural progenitor proliferation during cerebellar development (Leung et al. 2004), and activation of extracellular-regulated kinase (ERK), which plays a role in maintenance of pluripotency in normal human stem cells (Armstrong et al. 2006), in MB cells after treatment with NaB.

## Materials and Methods

### Cell Culture

Human Daoy (HTB186™) and D283 Med (HTB185™) MB cells were obtained from the American Type Culture Collection (ATCC, Rockville, USA). Cells were maintained in tissue culture flasks at 37 °C with humidified atmosphere and 5 % CO_2_. The culture medium, which was changed every 2/3 days, was prepared with DMEM (Gibco, Grand Island, USA), 1% penicillin and streptomycin (Gibco), supplemented with 10 % fetal bovine serum (Gibco), with the pH adjusted to 7.4.

### Treatments and Cell Viability

MB cells were seeded at 18.000 cells/well in 24 wells plates. After 24 h, cells were treated with the HDAC inhibitor NaB (Sigma-Aldrich, St. Louis, USA) alone or in combination with the MEK1/2 inhibitor U0126 (12.5 or 25 μM, Sigma-Aldrich). The NaB doses used in Daoy (0.25, 1.0, or 2.5 mM) and D283 (0.5, 2.5, or 5.0 mM) cells were chosen in agreement with a protocol previously described (Nör et al. 2013). Forty-eight hours after treatment, cells were trypsinized and counted in a Neubauer chamber with trypan blue (10 μl dye to each 10 μl cell suspension) for viability measurement.

### Histone H3 Acetylation

Cells treated for 48 h with NaB were lysed with a lysis solution buffer and acetylation of H3 was measured with PathScan^®^ Acetylated Histone H3 Sandwich enzyme-linked immunosorbent assay (ELISA) Kit (Cell Signaling Technology, Danvers, USA) according to the manufacturer’s instructions. Colorimetric signals were measured by spectrophotometric determination (OD450 nm) on Biochrom^®^ Anthos Zenyth 200 Microplate Reader.

### Gene Expression and Enrichment Analysis in MB Tumors from Patients

We analyzed data from 763 MB samples (Cavalli et al. 2017) to detect genes significantly correlated (*p* < 0.05) with both *CD133* or *BMI1* expression and to perform enrichment analysis of the correlated genes using the R2: Genomics Analysis and Visualization Platform (http://r2.amc.nl). KEGG results and Gene Ontology Biological Process pathways were checked again and plotted using the R software environment v.3.4.4 and the R package pathfindR v.1.2.1 (R Core Team, 2018; Ulgen et al., 2018), considering as significantly enriched pathways those with FDR < 0.05 in the function *run_pathfindR.* This function requires as input a dataset containing the gene identification (we used the HUGO symbol) the log fold change and corrected *p* value of their expression to perform the cut-off. Here, we replaced the fold change for the r (correlation) value and the p-value for the p-value of this correlation. Thus, we could indicate those negatively and positively correlated genes with either *BMI1* or *CD133.*

### Reverse Transcriptase Polymerase Chain Reaction (RT-qPCR)

The mRNA expression of stemness markers was analysed by RT-qPCR. Total RNA was extracted from MB cells using SV Total RNA Isolation System kit (Promega, Madison, USA), in accordance with the manufacturer’s instructions and quantified in NanoDrop (Thermo Fisher Scientific, Waltham, USA). The cDNA was obtained using GoScript Reverse System (Promega) according to the manufacturer’s instructions. The mRNA expression levels of target genes *(BMI1* and *CD133)* were quantified using PowerUp SYBR Green Master Mix (Thermo Fisher Scientific.) The primers used for RT-qPCR amplification were designed according to the corresponding GenBank and are shown in **Table 1**. Expression of β-actin was measured as an internal control.

**Table 1.**
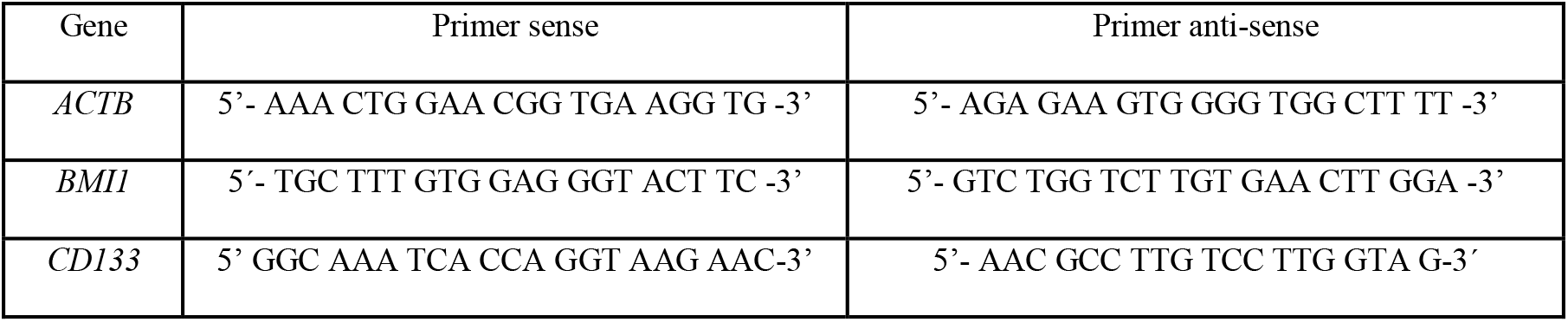
Forward and reverse primers used for RT-qPCR amplification

### Western Blot

Cells were lysed after NaB treatment with 1X Lysis Buffer (Cell Lysis Buffer 10X, Cell Signaling Technology), and protein was quantified using the Bradford protein assay (Thermo Scientific). For blotting, 20 μg of protein were separated by SDS-PAGE and transferred to a PVDF membrane. After 1h with blocking solution (5%BSA in TTBS), the membrane was incubated overnight at 4 °C with primary antibodies against BMI1 (1:1000; Cell Signaling), Phospho-p44/42 MAPK (pErk1/2, 1:2000; Cell Signaling), p44/42 MAPK (Erk1/2, 1:1000; Cell Signaling) and β-actin (1:2000; Santa Cruz Biotechnology) as protein control. Incubation of primary antibodies was followed by incubation with the secondary antibody (1:2000; Sigma Aldrich) for 2 h. Chemoluminescence was detected using ECL Western Blotting substrate (Pierce, Thermo Scientific) and analyzed using ImageQuant LAS 500 (GE Healthcare Life Sciences, Little Chalfont, UK). Immunodetection signal were analyzed using ImageJ (National Institutes of Health, Bethesda, USA).

### Neurosphere Formation

Neurosphere formation was used as an established experimental assay to evaluate cancer stem cell proliferation (Nör et al. 2013; Zanini et al. 2013). Daoy and D283 cells were dissociated with trypsin-EDTA into a single cell suspension and seeded at 500 cells/well in 24-well low-attachment plates. The cells were cultured in serum-free sphere-induction medium, containing DMEM/F12 supplemented with 20 ng/ml epidermal growth factor (EGF, Sigma-Aldrich), 20 ng/ml basic fibroblast growth factor (Sigma-Aldrich), B-27 supplement 1X (Gibco), N-2 supplement 0.5X (Gibco), 50 μg/ml bovine serum albumin (Sigma Aldrich), and antibiotics during 7 days as previously described (Nör et al. 2013). The ERK inhibitor U0126 was added in the first day after plating cells with the sphere-induction medium. Spheres photomicrographs were captured 7 days after treatment under an inverted phase microscope (Leica Microsystems, Mannheim Germany) at ×10 magnification.

### Statistics

Data are shown as mean ± standard deviation (SD). Statistical analysis was performed by one-way analysis of variance (ANOVA) followed by Tukey post-hoc tests for multiple comparisons. Experiments were replicated at least three times; *p* values under 0.05 were considered to indicate statistical significance. The GraphPad Prism 6 software (GraphPad Software, San Diego, USA) was used for analyses.

## Results

### HDAC Inhibition Impairs MB Cell Viability

Direct cell quantification after 48-h treatment with escalating doses of NaB confirmed a significant reduction in cell viability of both Daoy and D283 relative to vehicle. Consistently with a previous report (Nör et al. 2013), Daoy cells presented a somewhat higher sensibility to NaB than D283 cells. A dose of 2.5 mM reduced viability to 41.7% of control levels in Daoy cells *(p* < 0.0001), whereas the same dose decreased D283 cell viability to 71% of control levels *(p* < 0.0001). Lower doses of NaB, namely 0.25 mM in Daoy and 0.5 mM in D283 cells, did not significantly change cell numbers (**Fig. 1A**).

**Fig 1.**
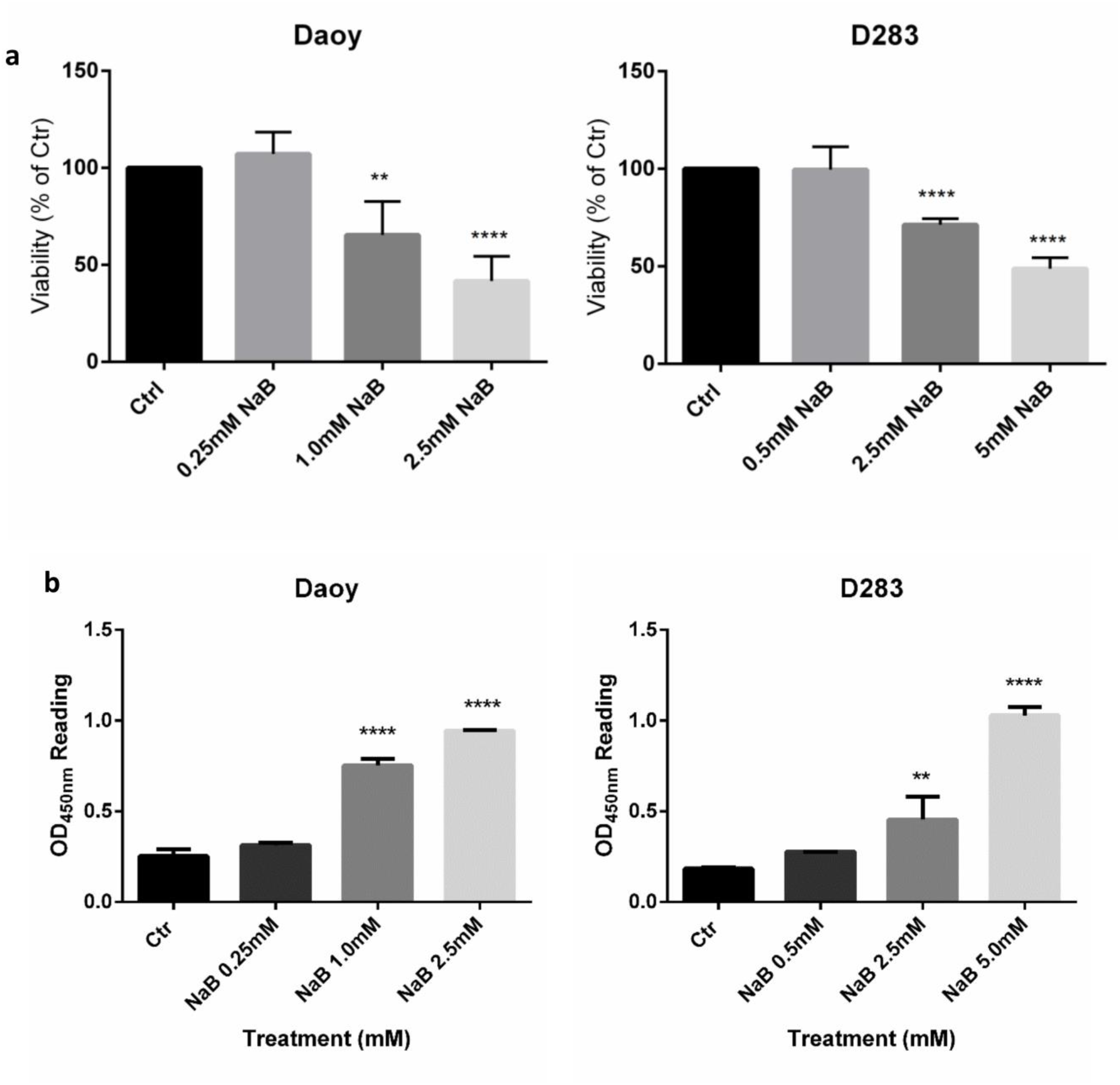
1 HDAC inhibition hinders the viability of MB cells. (A) Daoy and D283 cells were treated with different concentrations of NaB, and the number of living cells was quantified using trypan blue exclusion assays. (B) H3 acetylation was determined by ELISA. Results represent the mean ± SD of three independent experiments. ** *p* < 0.01; **** *p* < 0.0001 compared to controls (Ctr).

In order to evaluate a biochemical endpoint of efficacy, histone H3 acetylation was measured by ELISA in cells treated for 48 h. The effective doses of NaB (1.0 and 2.5 mM in Daoy cells, and 2.5 and 5.0 mM in D283 cells) induced 2.42-fold and 3.03-fold increases (Daoy, ps < 0.0001) and 2.5-fold and 5.7-fold increases (D283, *p* = 0.005 and *p* < 0.0001, respectively) in the levels of acetylated H3 compare to controls (**Fig. 1B**). Notably, the levels of acetylated H3 were inversely correlated to the effect on cell viability.

### HDAC Inhibition Reduces Expression of Stemness Markers and ERK Activation in MB

To elucidate whether NaB could suppress MB viability by modulating the stemness phenotype, expression of *BMI1* and *CD133* was analyzed after 48 h of NaB treatment. Both marks are expressed in all molecular subgroups of MB as observed in Cavalli et al. (2017) samples (**Fig. 2A**). Experiments using RT-qPCR showed a reduction in mRNA levels of stemness markers *BMI1* and *CD133.* In Daoy cells, mRNA expression of *BMI1* was reduced in cells treated with the higher dose of NaB (2.5 mM; *p* = 0.0007), whereas all doses of NaB (0.25, 1.0, and 2.5 mM) reduced *CD133* expression compared to controls (0.25 mM, *p* = 0.018; 1 mM, *p* = 0.01; 2.5 mM, *p* = 0.0006). In D283 cells, both doses that were effective when used in cell counting experiments (2.5 and 5 mM) reduced *BMI1* expression (2.5 mM, p = 0.0017; 5 mM, *p* < 0.0001), and *CD133* expression was reduced by all doses (0.5 mM, *p* = 0.0198; 2.5 mM and 5 mM, *p* < 0.0001) (**Fig. 2B**).

**Fig 2.**
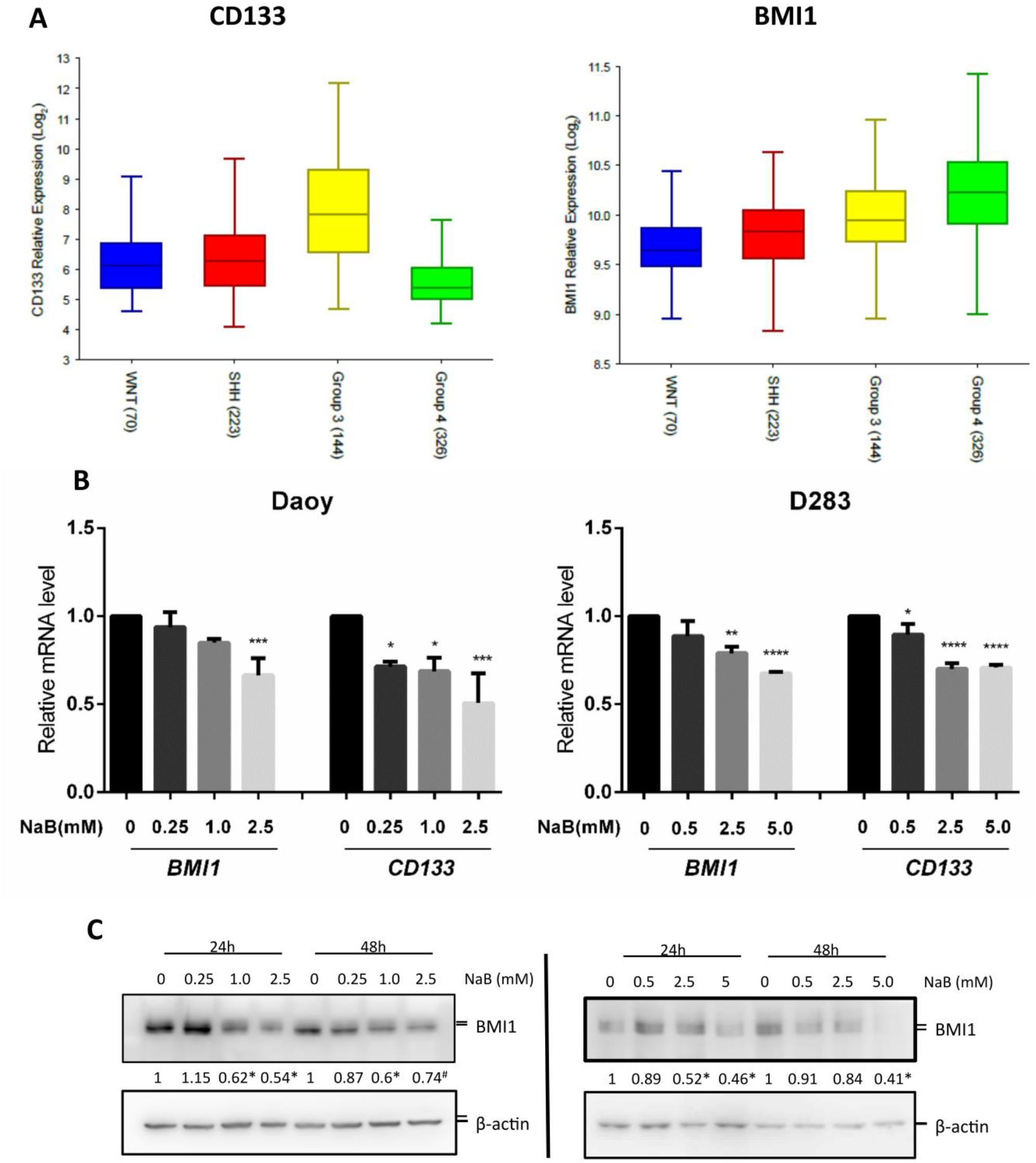
HDAC inhibition reduces stemness in MB cells. (A) Representative boxplot of *CD133* and *BMI1* expression in the medulloblastoma molecular subgroups from Cavalli et al. (30). (B) Relative mRNA levels of stemness markers were determined in control and NaB-treated Daoy and D283 MB cells by RT-qPCR using primers specific for *BMI1* and *CD133* as described in Materials and Methods. (C) Daoy and D283 cells were treated with different concentrations of NaB for 24 or 48h, cells were then harvested and the expression of BMI1 and β-actin (loading control) was determined by Western blot analysis. The relative expression of BMI1 (24 or 48 h), were determined by densitometric analysis of signals normalized for the corresponding β-actin signal. The analysis was performed using Image J software (NIH, Bethesda, MD). Results represent the mean ± SD of three independent experiments; * *p* < 0.05; ** *p* < 0.01; *** *p* < 0.001; **** *p* < 0.0001 compared to controls (Ctr).

The reduction in protein levels of BMI1 was confirmed after 24 h of NaB treatment in both cell lines at any dose capable of significantly impairing cell viability. After a 48-h treatment, Daoy cells maintained a consistent reduction in BMI1 protein levels compared to controls, and a significant reduction was found in D283 cells treated with the highest dose (5 mM) compared to controls *(p* = 0.0023) (**Fig. 2C**).

MAPK/ERK signaling regulates normal CNS development. The enrichment analysis of genes significantly correlated with *CD133* or *BMI1* expression in MB tumor samples from patients showed MAPK signaling or associated pathways (such as neurotrophin and Erb signaling) as the most enriched process (**Fig. 3A, Supplementary Table S1, S2, Supplementary Fig. S1, S2**). To investigate the status of the MAPK/ERK pathway in the MB cells after treatment with NaB, Western blotting was performed. NaB at any dose led to a decrease in ERK pathway activation measured by ERK phosphorylation in both cell lines after 48 h. Even in non-effective doses (0.25 mM in Daoy and 0.5 mM in D283 cells), a reduction of about 20% was observed, whereas doses that significantly affect cell viability reduced ERK activation in both cell lines (*ps* < 0.0001, **Fig. 3B, 3C**).

**Fig 3.**
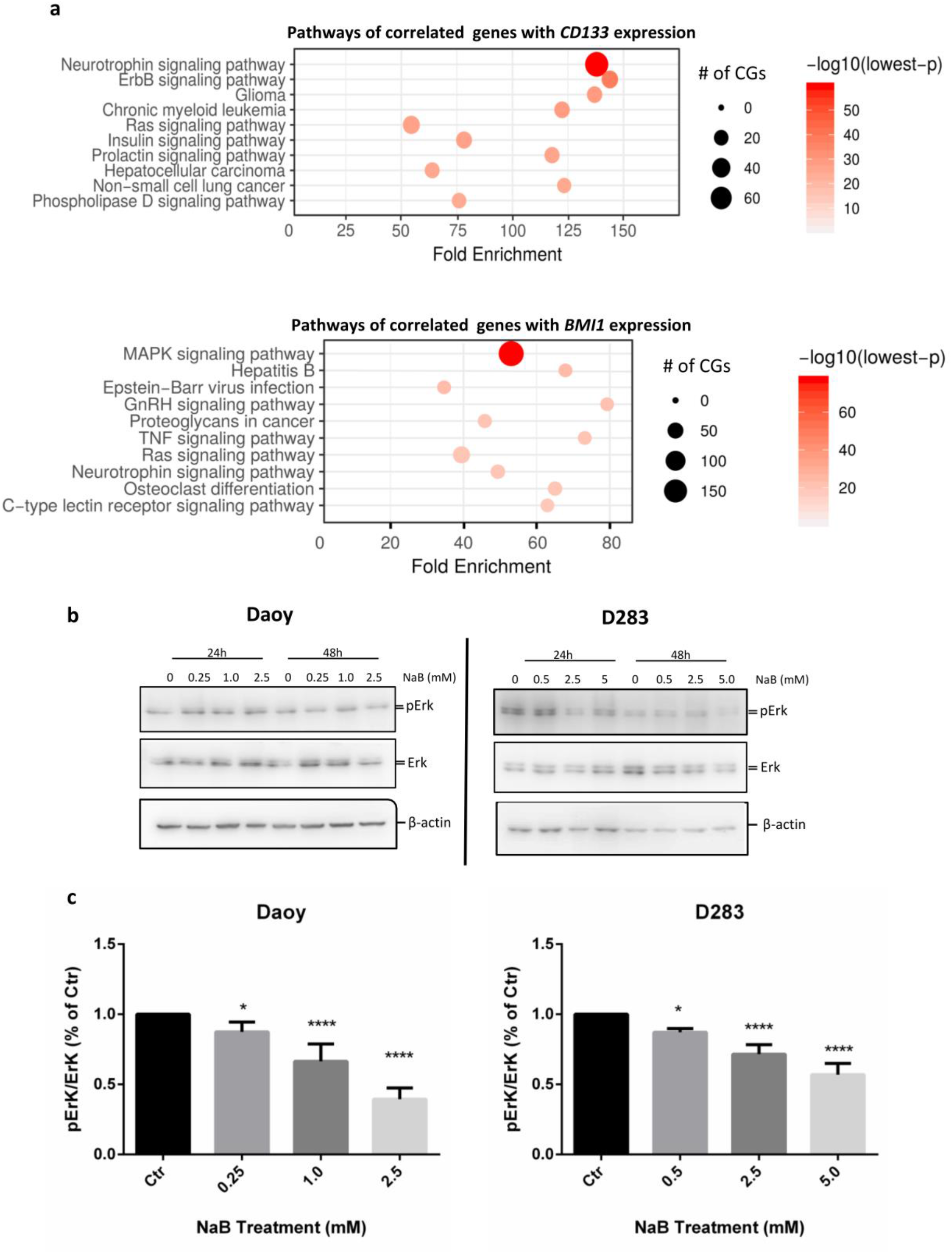
HDAC inhibition reduces ERK activation in MB cells. (A) Balloon plot of the 10 most representative KEGG pathways of *CD133* or *BMI1* correlated genes. # of CGs: number of correlated genes. Large circles indicate higher number of correlated genes. A darker color indicates a more significant *p* value. (B) Daoy and D283 cells were treated with different concentrations of NaB for 24 or 48 h, cells were then harvested and the expression ERK, pERK and β-actin (loading control) was determined by Western blot analysis. The relative expression of ERK and p-ERK (48 h) were determined by densitometric analysis of signals normalized for the corresponding β-actin signal. The analysis was performed using Image J software (NIH, Bethesda, MD). Results represent the mean ± SD of three independent experiments; *\*p* < 0.05; \*\**p* < 0.01; \*\*\**p* < 0.001; **** *p* < 0.0001 compared to controls (Ctr).

### HDAC and MAPK/ERK Inhibition Cooperate to Reduce MB Growth

To determine the effect of ERK inhibition on stemness markers in MB cells, mRNA expression of *BMI1* and *CD133* was measured after 48 h of treatment with the MEK1/2 inhibitor U0126 (12.5, or 25 μM). *CD133* expression decreased significantly after treatment in both cell lines (Daoy, *ps* = 0.0001; D283, *p* = 0.02 and *p* < 0.0001). Reductions in *BMI1* were observed in both cell lines (Daoy, 12.5 μM, *p* = 0.0094; 25 μM, *p* = 0.0016; D283, 25 μM, *p* = 0.0436) (**Fig. 4A**).

**Fig 4.**
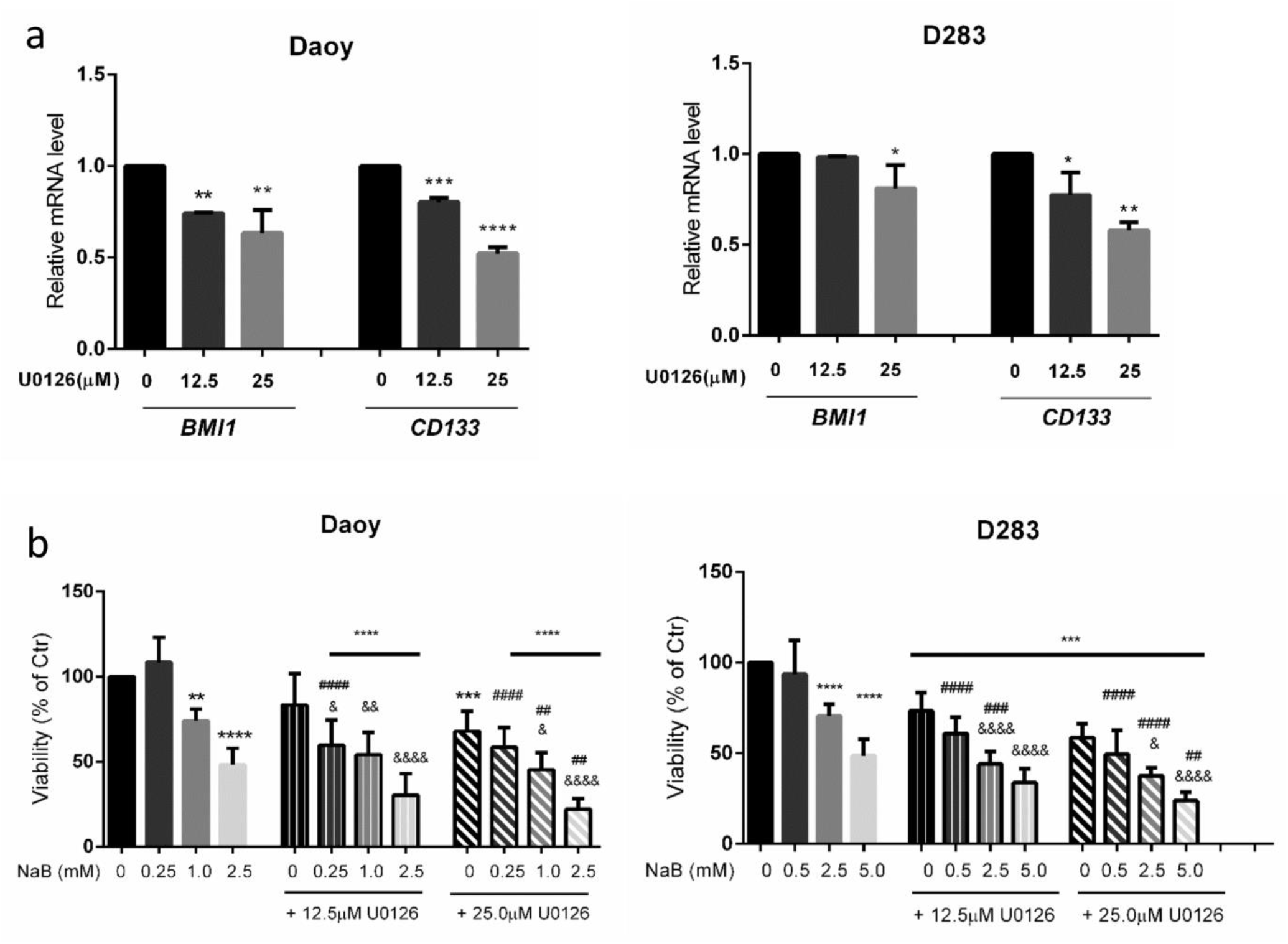
Inhibition of HDAC and ERK impairs MB cell proliferation. (A) Relative mRNA levels of stemness markers were determined in control and U0126-treated Daoy and D283 MB cells by RT-qPCR, using primers specific for *BMI1* and *CD133.* (B) Daoy and D283 cells were treated with different concentrations of NaB alone or in combination with the ERK inhibitor U0126, and the number of living cells were quantified using trypan blue exclusion assays. Results represent the mean ± SD of three independent experiments; \**p* < 0.5; \*\**p* < 0.01; \*\*\**p* < 0.001; \*\*\*\**p* < 0.0001 compared to controls (Ctr); #p < 0.05; ##*p* < 0.01; ###*p* < 0.001; ####*p* < 0.0001 compared to respective NaB dose; & *p* < 0.05; &&&& *p* < 0.0001 compared to respective U0126 dose.

Given that there is a possible relationship between the stemness phenotype and cell viability, cell counting after treatment with U0126 with or without NaB was performed. U0126 (25 μM) significantly reduced MB cell viability (Daoy, *p* = 0.0008; D283, *p* < 0.0001). Combined treatment of the highest dose of U0126 (25 μM) plus NaB (1.0 or 2.5 mM in Daoy and 2.5 or 5 mM in D283) produced more robust inhibition of cell proliferation than either inhibitor alone at the same doses in both cell lines. (**Fig. 4B**).

### MAPK/ERK Inhibition Hinders the Expansion of Putative MB CSCs

A neurosphere assay was performed to investigate the formation of MB CSCs. First, the mRNA expression of *CD133* and *BMI1* was measured to confirm enrichment of stemness after sphere formation. Expression of both markers was increased in neurospheres compared to monolayer cultures. After 7 days of culturing cells in appropriated medium for expansion of stem cells, *BMI1* expression was increased 1.68-fold in Daoy *(p* = 0.0006) and 2.0-fold in D283 *(p* = 0.0028) cells, while *CD133* expression increased 2.89-fold in Daoy *(p* = 0.0005) and to 2.36-fold in D283 *(p* < 0.0001) cells in comparison to controls (**Fig. 5A**). After MAPK/ERK inhibition (12. μM or 25 μM), cells were unable to form neurospheres (**Fig. 5B**). These results suggest an essential role of ERK signalling in MB stem cell formation.

**Fig 5.**
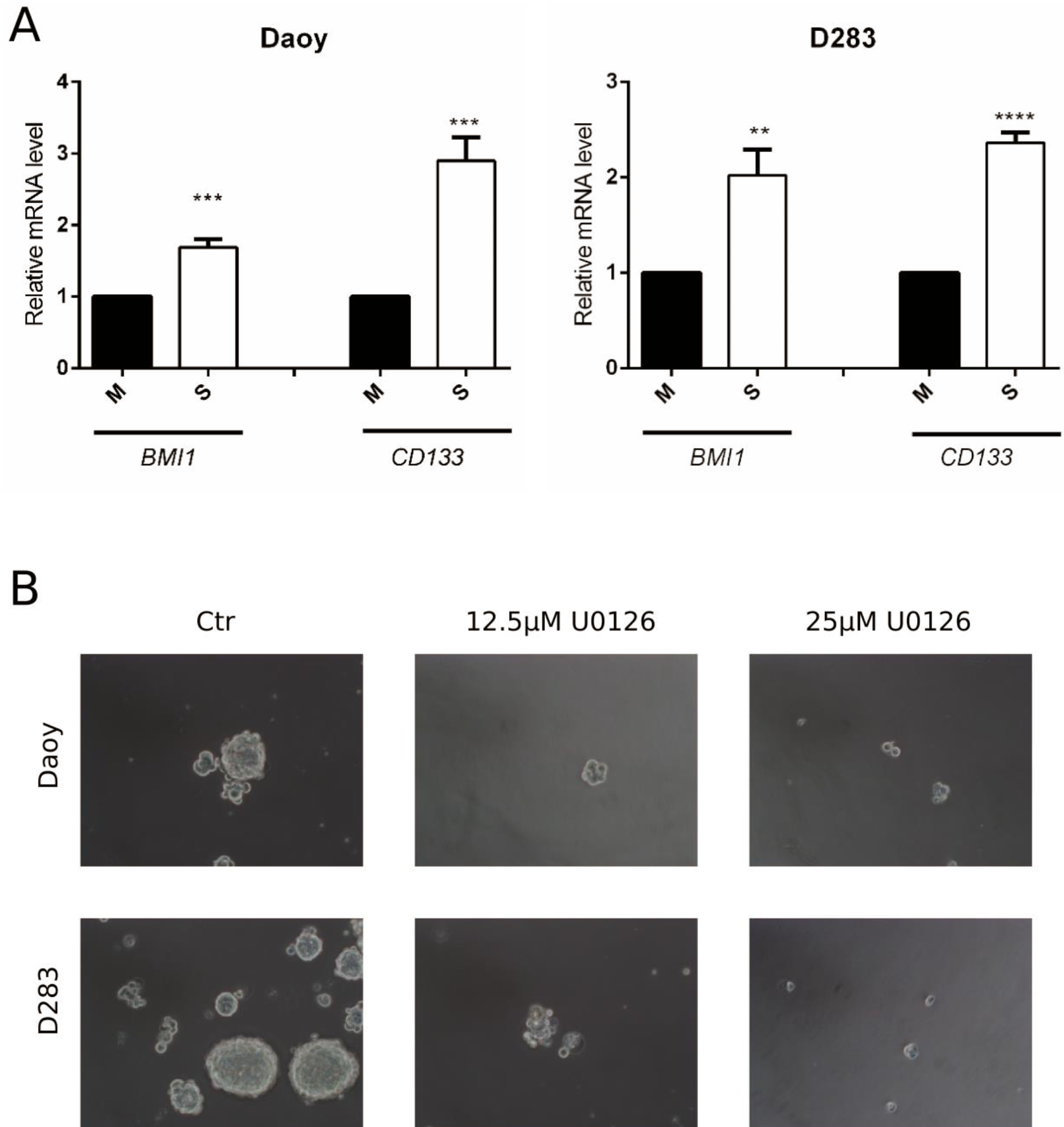
ERK inhibition hinders MB stem cell formation. (A) Daoy and D283 cells were cultured in medium specific for stem cell expansion during 7 days for enrichment of neurospheres. Relative mRNA levels of stemness markers were determined in monolayer (M) and sphere (S) cultures by RT-qPCR using primers specific for *BMI1* and *CD133* as described in Materials and Methods. (B) Representative photomicrographs of neurospheres after 7 days of induction in the absence (Ctr) or presence of the ERK inhibitor U0126 (12. μM or 25 μM) in the first day of induction. Results represent the mean ± SD of three independent experiments; \*\**p* < 0.01; \*\*\**p* < 0.001; **** *p* < 0.0001 compared to controls.

## Discussion

MB is a pediatric brain tumor featuring a high frequency of mutations in epigenetic modifier genes (Robinson et al. 2012; Huether et al. 2014). Deregulation in epigenetic programming has been linked to tumorigenesis, possibly through induction of a stemness phenotype (Hadjimichael et al. 2015). Histone acetylation status is related to cancer stemness regulation, and HDAC inhibition constitutes an important therapeutic strategy to induce differentiation in CSCs (Liu et al. 2017). Abnormal epigenetic regulation related to histone modifications have been recently reported in SHH MB (Robinson et al. 2019).

We have previously shown responsiveness of human MB cells and putative MB CSCs to NaB (Nör et al. 2013). Here, we used the same compound to investigate the influence of HDAC inhibition on maintenance of a stemness phenotype of MB. We confirmed that the effect of NaB is associated with its capability to increase histone acetylation. Based on results from our analyses of gene expression in MB samples from patients, we have also shown a reduction in markers of stemness and inhibition of MAPK/ERK signalling, and that HDAC and MEK1/2 inhibitors can cooperate to reduce viability in MB cells and, as with HDACi, MAPK/ERK inhibition hinders neurosphere formation. The cell lines we used have been characterized to represent distinct MB molecular subgroups: D283 cells display *MYC* amplification and show features of Groups 3 and 4 MB, whereas Daoy cells are *TP53*-mutated and represent SHH MB (Ivanov et al. 2016; Higdon et al. 2017; Thompson et al. 2017; Bonfim-Silva et al. 2019).

Expression of *CD133* and *BMI1* was significantly decreased after NaB treatment. CD133, also called Prominin-1, is the most used marker to isolate and to identify brain tumor CSCs (Schonberg et al. 2014). BMI1 is a polycomb group protein overexpressed in MB (Leung et al., 2004) and present in CSC (Manoranjan et al. 2013). Patients with metastatic MB show higher levels of *CD133* and a worse prognostic when compared with non-metastatic patients (Raso et al. 2011). *In vitro* characterization of CD133-positive cells demonstrates that they present increased migration and invasion capabilities and higher expression of stemness-associated genes compared to the parental cell line. In addition, CD133-positive, but not CD133-negative cells, are invasive *in vivo* (Chen et al. 2010; Chang et al. 2012). These functional features of CD133-positive cells are associated with a stemness genetic signature. The volume of xenografted tumors is reduced by treatments that inhibit the expression of stemness-related genes such as *BMI1, SOX2, Nestin, Nanog,* and *Oct4* (Zakrzewska et al. 2011; Chang et al. 2012). BMI-1, another marker of poor outcomes in MB (Zakrzewska et al. 2011), is also involved in tumor invasion, as indicated by evidence that BMI1 knockdown in MB cells decreases infiltration of xenotransplants (Wang et al. 2012).

We observed a decrease in phosphorylated ERK (pERK) in MB cells after NaB treatment. Components of the ERK pathway are upregulated in metastatic MB, and pharmacological inhibition of ERK decreases migration of MB cells (MacDonald et al. 2001). In carcinoma and sarcoma cell lines, ERK reduces the stem-like characteristics of tumor cells (Tabu et al. 2010). Increased expression of genes in the MAPK/ERK pathway is related to poor survival outcome in MB patients (Park et al. 2019). In our study, ERK inhibition reduced the expression level of stemness markers and enhanced the antiproliferative effect of NaB. In agreement with our results, Ozaki and colleagues have shown an increase in HDACi-induced cell death after specific inhibition of the ERK pathway by the generation of reactive oxygen species (Ozaki et al. 2006; 2010). Although our study has not evaluated the mechanisms mediating functional interactions between HDAC and ERK inhibitors, our findings suggest that both types of compounds are involved in stemness modulation.

The influence of ERK activity in regulating the stemness phenotype was showed in a neurosphere model (Ishiguro et al. 2017). Expansion of neurospheres resulted in an enrichment of the stemness phenotype in cell cultures, as indicated by increased levels of *CD133* and *BMI1* content in comparison with a monolayer culture. We found that pharmacological inhibition of ERK impairs MB neurosphere formation. The role of ERK signaling was already described in other tumor stem-like cells (Sunayama et al. 2010; Ciccarelli et al. 2016). In embryonal rhabdomyosarcoma stem-like cells, the inhibition of ERK decrease sphere formation, induce myogenic differentiation, and reduce tumorigenicity *in vivo* (Ciccarelli et al. 2016). In glioblastoma CSCs, ERK inhibition also limited the capacity to form tumors in nude mice and promote cell differentiation. Consistently with our results, ERK inhibition of glioblastoma CSCs reduces the expression of stemness markers, including *BMI1* (Sunayama et al. 2010).

In MB, Chow et al. (2014) showed differential ERK signalling activation and neurosphere formation capability in MB cells from spontaneous tumors obtained from Patched (Ptch)+/− mice. Tumors that were able to form neurospheres independently of the presence of growth factors (EGF and basic fibroblast growth factor) had a higher activation of the ERK pathway compared to those tumors dependent on growth factors or not capable of forming spheres. Interestingly, tumors derived from growth factors independent neurospheres cells presented higher tumorigenicity potential and lethality.

It should be pointed out that our study has several limitations, including the lack of in vivo experiments and patient-derived primary cell cultures, and the typical limitations related to cell line cultures, such as the possibility of genetic and phenotypic drift, artificial culture environment, and the lack of cell matrix interactions and immune regulation.

In summary, our results provide evidence indicating that HDAC inhibition can modulate the stemness phenotype, by reducing the expression of stemness markers *CD133* and *BMI1,* and decreasing ERK activity, in MB. The findings indicate for the first time that combined inhibition of HDAC and ERK signaling was more effective in reducing MB cell viability than single-drug treatments. This is also the first report to indicate that ERK inhibition impairs MB CSC formation/survival. We suggest the combination of HDAC and ERK inhibitors as an experimental therapeutic approach in MB, which should be further investigated with the use of additional experimental approaches such as *in vivo* models.

## Acknowledgements

This manuscript has been released as a pre-print at bioRxiv: https://www.biorxiv.org/content/10.1101/521393v1. This research was supported by the National Council for Scientific and Technological Development (CNPq; grant numbers 303276/2013-4 and 409287/2016-4 to R.R., grant number 201001/2014-4 to C.N., and graduate fellowship to M.C.J.); PRONON/Ministry of Health, Brazil (number 25000.162.034/2014-21); the Children’s Cancer Institute (ICI); the Rio Grande do Sul State Research Foundation (FAPERGS; grant number 17/2551-0001 071-0 to R.R.); the Coordination for the Improvement of Higher Education Personnel (CAPES); and the Clinical Hospital institutional research fund (FIPE/HCPA). T.F. is supported by Programa Nacional de Pós-Doutorado (PNPD, CAPES/HCPA, grant number: 88887.160608/2017-00). C.N. is also supported by the William Donald Nash fellowship from the Brain Tumour Foundation of Canada. V.R. is supported by operating funds from the Canadian Institutes for Health Research, the American Brain Tumor Association, and the Brain Tumour Foundation of Canada. M.D.T. is supported by the National Institutes of Health, the Pediatric Brain Tumour Foundation, the Terry Fox Research Institute, the Canadian Institutes of Health Research, the Cure Search Foundation, b.r.a.i.n.child, Meagan’s Walk, Genome Canada, Genome BC, the Ontario Institute for Cancer Research, and the Canadian Cancer Society Research Institute.

## Supplementary material

The Supplementary Material for this article can be found online.

