## Supplemental information for "HDAC and MAPK/ERK Inhibitors Cooperate to Reduce Viability and Stemness in Medulloblastoma"

**Supplementary Fig S1** Balloon plot of significantly enriched KEGG pathways of CD133 correlated genes. # of CGs: number of correlated genes. Large circles indicate higher number of correlated genes. A darker color indicates a more significant *p* value.

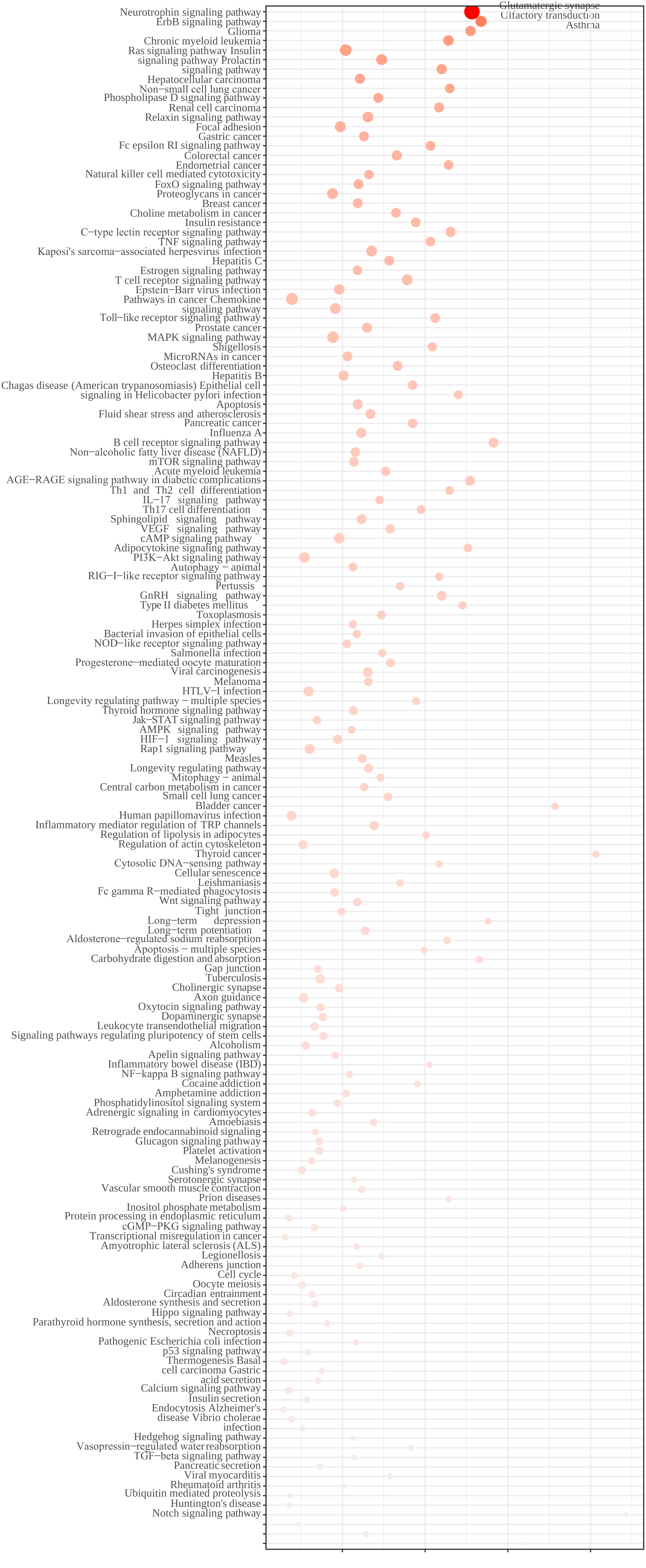
Glutamatergic synapse
Olfactory transduction
Asthma
Neurotrophin signaling pathway
ErbB signaling pathway
Glioma
Chronic myeloid leukemia
Ras signaling pathway Insulin
signaling pathway Prolactin
signaling pathway
Hepatocellular carcinoma
Non−small cell lung cancer
Phospholipase D signaling pathway
Renal cell carcinoma
Relaxin signaling pathway
Focal adhesion
Gastric cancer
Fc epsilon RI signaling pathway
Colorectal cancer
Endometrial cancer
Natural killer cell mediated cytotoxicity
FoxO signaling pathway
Proteoglycans in cancer
Breast cancer
Choline metabolism in cancer
Insulin resistance
C−type lectin receptor signaling pathway
TNF signaling pathway
Kaposi's sarcoma−associated herpesvirus infection
Hepatitis C
Estrogen signaling pathway
T cell receptor signaling pathway
Epstein−Barr virus infection
Pathways in cancer Chemokine
signaling pathway
Toll−like receptor signaling pathway
Prostate cancer
MAPK signaling pathway
Shigellosis
MicroRNAs in cancer
Osteoclast differentiation
Hepatitis B
Chagas disease (American trypanosomiasis) Epithelial cell
signaling in Helicobacter pylori infection
Apoptosis
Fluid shear stress and atherosclerosis
Pancreatic cancer
Influenza A
B cell receptor signaling pathway
Non−alcoholic fatty liver disease (NAFLD)
mTOR signaling pathway
Acute myeloid leukemia
AGE−RAGE signaling pathway in diabetic complications
Th1 and Th2 cell differentiation
IL−17 signaling pathway
Th17 cell differentiation
Sphingolipid signaling pathway
VEGF signaling pathway
cAMP signaling pathway
Adipocytokine signaling pathway
PI3K−Akt signaling pathway
Autophagy − animal
RIG−I−like receptor signaling pathway
Pertussis
GnRH signaling pathway
Type II diabetes mellitus
Toxoplasmosis
Herpes simplex infection
Bacterial invasion of epithelial cells
NOD−like receptor signaling pathway
Salmonella infection
Progesterone−mediated oocyte maturation
Viral carcinogenesis
Melanoma
HTLV−I infection
Longevity regulating pathway − multiple species
Thyroid hormone signaling pathway
Jak−STAT signaling pathway
AMPK signaling pathway
HIF−1 signaling pathway
Rap1 signaling pathway
Measles
Longevity regulating pathway
Mitophagy − animal
Central carbon metabolism in cancer
Small cell lung cancer
Bladder cancer
Human papillomavirus infection
Inflammatory mediator regulation of TRP channels
Regulation of lipolysis in adipocytes
Regulation of actin cytoskeleton
Thyroid cancer
Cytosolic DNA−sensing pathway
Cellular senescence
Leishmaniasis
Fc gamma R−mediated phagocytosis
Wnt signaling pathway
Tight junction
Long−term depression
Long−term potentiation
Aldosterone−regulated sodium reabsorption
Apoptosis − multiple species
Carbohydrate digestion and absorption
Gap junction
Tuberculosis
Cholinergic synapse
Axon guidance
Oxytocin signaling pathway
Dopaminergic synapse
Leukocyte transendothelial migration
Signaling pathways regulating pluripotency of stem cells
Alcoholism
Apelin signaling pathway
Inflammatory bowel disease (IBD)
NF−kappa B signaling pathway
Cocaine addiction
Amphetamine addiction
Phosphatidylinositol signaling system
Adrenergic signaling in cardiomyocytes
Amoebiasis
Retrograde endocannabinoid signaling
Glucagon signaling pathway
Platelet activation
Melanogenesis
Cushing's syndrome
Serotonergic synapse
Vascular smooth muscle contraction
Prion diseases
Inositol phosphate metabolism
Protein processing in endoplasmic reticulum
cGMP−PKG signaling pathway
Transcriptional misregulation in cancer
Amyotrophic lateral sclerosis (ALS)
Legionellosis
Adherens junction
Cell cycle
Oocyte meiosis
Circadian entrainment
Aldosterone synthesis and secretion
Hippo signaling pathway
Parathyroid hormone synthesis, secretion and action
Necroptosis
Pathogenic Escherichia coli infection
p53 signaling pathway
Thermogenesis Basal
cell carcinoma Gastric
acid secretion
Calcium signaling pathway
Insulin secretion
Endocytosis Alzheimer's
disease Vibrio cholerae
infection
Hedgehog signaling pathway
Vasopressin−regulated water reabsorption
TGF−beta signaling pathway
Pancreatic secretion
Viral myocarditis
Rheumatoid arthritis
Ubiquitin mediated proteolysis
Huntington's disease
Notch signaling pathway

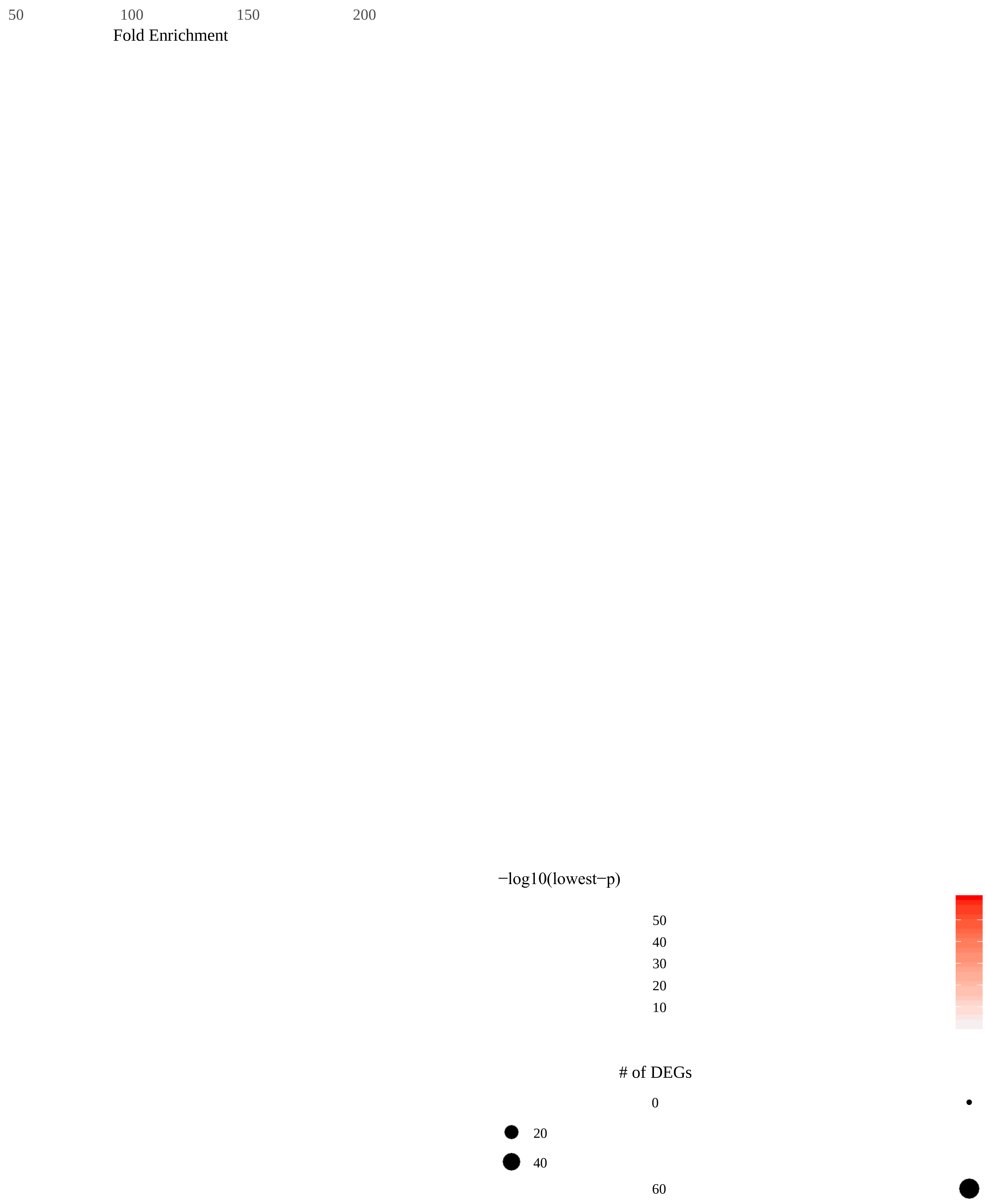
50
100
150
200
Fold Enrichment
−log10(lowest−p)
50
40
30
20
10
### of DEGs
0
20
40
60

**Supplementary Fig S2** Balloon plot of significantly enriched KEGG pathways of BMI1 correlated genes. # of CGs: number of correlated genes. Large circles indicate higher number of correlated genes. A darker color indicates a more significant *p* value.

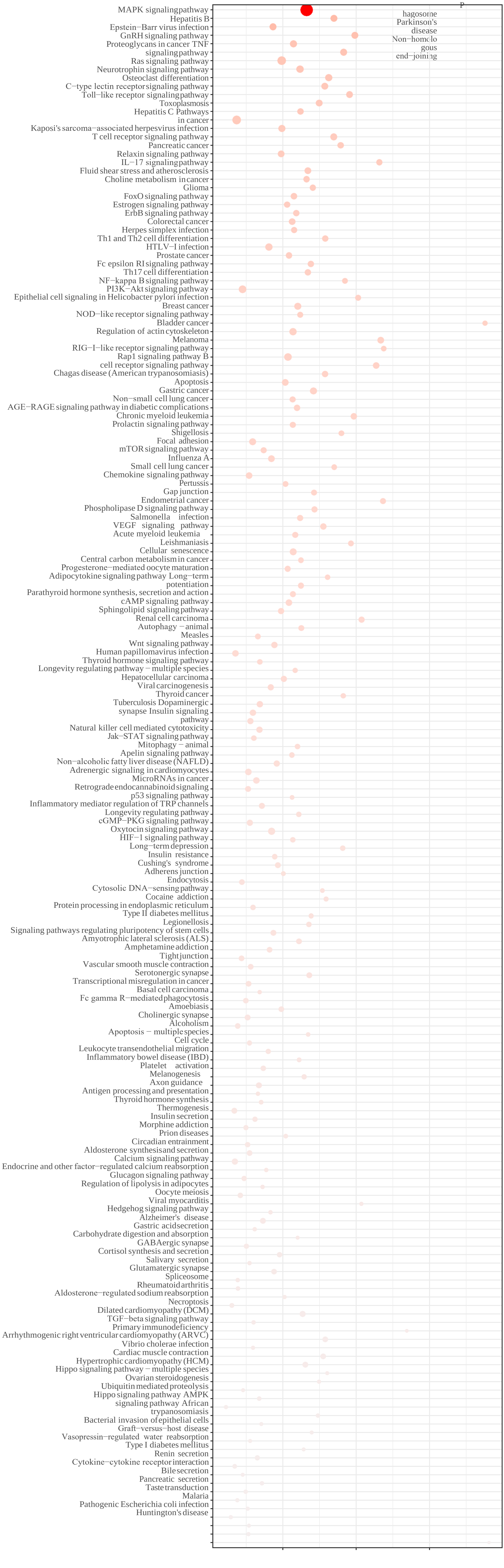

MAPK signalingpathway
Hepatitis B
Epstein−Barr virus infection
GnRH signaling pathway
Proteoglycans in cancer TNF
signalingpathway
Ras signaling pathway
Neurotrophin signaling pathway
Osteoclast differentiation
C−type lectin receptorsignaling pathway
Toll−like receptor signalingpathway
Toxoplasmosis
Hepatitis C Pathways
in cancer
Kaposi's sarcoma−associated herpesvirus infection
T cell receptor signaling pathway
Pancreatic cancer
Relaxin signaling pathway
IL−17 signalingpathway
Fluid shear stress and atherosclerosis
Choline metabolism in cancer
Glioma
FoxO signaling pathway
Estrogen signaling pathway
ErbB signaling pathway
Colorectal cancer
Herpes simplex infection
Th1 and Th2 cell differentiation
HTLV−I infection
Prostate cancer
Fc epsilon RI signaling pathway
Th17 cell differentiation
NF−kappa B signaling pathway
PI3K−Akt signaling pathway
Epithelial cell signaling in Helicobacter pylori infection
Breast cancer
NOD−like receptor signaling pathway
Bladder cancer
Regulation of actin cytoskeleton
Melanoma
RIG−I−like receptor signaling pathway
Rap1 signaling pathway B
cell receptor signaling pathway
Chagas disease (American trypanosomiasis)
Apoptosis
Gastric cancer
Non−small cell lung cancer
AGE−RAGE signaling pathway in diabetic complications
Chronic myeloid leukemia
Prolactin signaling pathway
Shigellosis
Focal adhesion
mTOR signaling pathway
Influenza A
Small cell lung cancer
Chemokine signaling pathway
Pertussis
Gap junction
Endometrial cancer
Phospholipase D signaling pathway
Salmonella infection
VEGF signaling pathway
Acute myeloid leukemia
Leishmaniasis
Cellular senescence
Central carbon metabolism in cancer
Progesterone−mediated oocyte maturation
Adipocytokine signaling pathway Long−term
potentiation
Parathyroid hormone synthesis, secretion and action
cAMP signaling pathway
Sphingolipid signaling pathway
Renal cell carcinoma
Autophagy − animal
Measles
Wnt signaling pathway
Human papillomavirus infection
Thyroid hormone signaling pathway
Longevity regulating pathway − multiple species
Hepatocellular carcinoma
Viral carcinogenesis
Thyroid cancer
Tuberculosis Dopaminergic
synapse Insulin signaling
pathway
Natural killer cell mediated cytotoxicity
Jak−STAT signaling pathway
Mitophagy − animal
Apelin signaling pathway
Non−alcoholic fatty liver disease (NAFLD)
Adrenergic signaling in cardiomyocytes
MicroRNAs in cancer
Retrograde endocannabinoid signaling
p53 signaling pathway
Inflammatory mediator regulation of TRP channels
Longevity regulating pathway
cGMP−PKG signaling pathway
Oxytocin signaling pathway
HIF−1 signaling pathway
Long−term depression
Insulin resistance
Cushing's syndrome
Adherens junction
Endocytosis
Cytosolic DNA−sensing pathway
Cocaine addiction
Protein processing in endoplasmic reticulum
Type II diabetes mellitus
Legionellosis
Signaling pathways regulating pluripotency of stem cells
Amyotrophic lateral sclerosis (ALS)
Amphetamine addiction
Tight junction
Vascular smooth muscle contraction
Serotonergic synapse
Transcriptional misregulation in cancer
Basal cell carcinoma
Fc gamma R−mediated phagocytosis
Amoebiasis
Cholinergic synapse
Alcoholism
Apoptosis − multiple species
Cell cycle
Leukocyte transendothelial migration
Inflammatory bowel disease (IBD)
Platelet activation
Melanogenesis
Axon guidance
Antigen processing and presentation
Thyroid hormone synthesis
Thermogenesis
Insulin secretion
Morphine addiction
Prion diseases
Circadian entrainment
Aldosterone synthesis and secretion
Calcium signaling pathway
Endocrine and other factor−regulated calcium reabsorption
Glucagon signaling pathway
Regulation of lipolysis in adipocytes
Oocyte meiosis
Viral myocarditis
Hedgehog signaling pathway
Alzheimer's disease
Gastric acid secretion
Carbohydrate digestion and absorption
GABAergic synapse
Cortisol synthesis and secretion
Salivary secretion
Glutamatergic synapse
Spliceosome
Rheumatoid arthritis
Aldosterone−regulated sodium reabsorption
Necroptosis
Dilated cardiomyopathy (DCM)
TGF−beta signaling pathway
Primary immunodeficiency
Arrhythmogenic right ventricular cardiomyopathy (ARVC)
Vibrio cholerae infection
Cardiac muscle contraction
Hypertrophic cardiomyopathy (HCM)
Hippo signaling pathway − multiple species
Ovarian steroidogenesis
Ubiquitin mediated proteolysis
Hippo signaling pathway AMPK
signaling pathway African
trypanosomiasis
Bacterial invasion of epithelial cells
Graft−versus−host disease
Vasopressin−regulated water reabsorption
Type I diabetes mellitus
Renin secretion
Cytokine−cytokine receptor interaction
Bile secretion
Pancreatic secretion
Taste transduction
Malaria
Pathogenic Escherichia coli infection
Huntington's disease
P
hagosome
Parkinson's
disease
Non−homolo
gous
end−joining

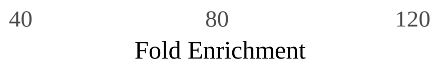
40
80
120
Fold Enrichment

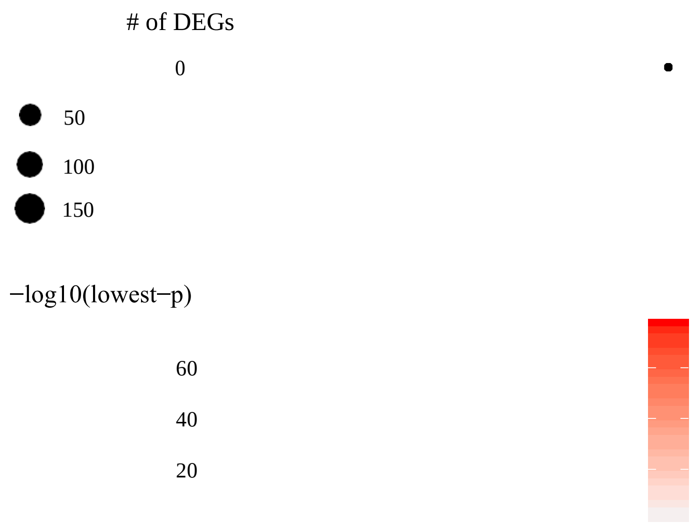
### of DEGs
0
50
100
150
−log10(lowest−p)
60
40
20

**Supplementary Table S1** Enrichment analysis (KEGG pathways) of CD133-correlated genes.

| ID | Pathway | Fold_Enrichment | occurrence | lowest_p | highest_p | Up_regulated | Down_regulated |
|---|---|---|---|---|---|---|---|
| hsa04722 | Neurotrophin signaling pathway | 128,3859649 | 10 | 1,96279E-60 | 1,87915E-49 | PIK3CB, PIK3R1, PLCG2, SHC1, SHC2, HRAS, MAP2K2, RPS6KA1, CRK, FOXO3, NFKBIA, NFKBIB, ATF4, SORT1, CAMK4, CAMK2A, CAMK2D, CAMK2G, MATK, RHOA, TP53, MAP3K1, JUN, IRS1, MAPKAPK2, MAP2K5, ARHGDIG, SH2B3, MAP2K7, PRDM4, RIPK2, IRAK4, YWHAE, IKBKB, PSEN2, BAX | NTRK3, PIK3CA, PIK3R2, PLCG1, SHC4, GRB2, NGF, NGFR, AKT1, AKT3, BRAF, SOS1, SOS2, MAPK3, RAPGEF1, RAP1B, PTPN11, GAB1, GSK3B, NFKBIE, NFKB1, RELA, FRS2, PDPK1, KIDINS220, RPS6KA5, MAPK8, MAPK9, MAPK10, MAPK11, MAPK7, ARHGDIA, SH2B2, SH2B1, ABL1, CDC42, RAC1, TP73, MAGED1, IRAK2, CALM3 |
| hsa04012 | ErbB signaling pathway | 133,8454504 | 10 | 6,08872E-39 | 7,25968E-33 | MAP2K7, CRK, SHC1, SHC2, CAMK2A, CAMK2D, CAMK2G, PLCG2, JUN, PIK3CB, PIK3R1, MAP2K2, HRAS | GRB2, GSK3B, GAB1, MAPK8, MAPK9, MAPK10, SOS1, SOS2, SHC4, PLCG1, ABL1, PIK3CA, PIK3R2, AKT1, AKT3, MAPK3, BRAF |

| | | | | | | | |
|---|---|---|---|---|---|---|---|
| hsa05214 | Glioma | 127,546841 | 10 | 8,30394E-30 | 2,35549E-27 | CAMK2A, CAMK2D, CAMK2G, SHC1, SHC2, MAP2K2, PIK3CB, PIK3R1, TP53, PLCG2, HRAS, BAX | CALM3, SHC4, GRB2, SOS1, SOS2, BRAF, MAPK3, PIK3CA, PIK3R2, AKT1, AKT3, PLCG1 |
| hsa05220 | Chronic myeloid leukemia | 114,1208577 | 10 | 2,09995E-29 | 4,79749E-26 | TP53, NFKBIA, IKBKB, SHC1, SHC2, CRK, PIK3CB, PIK3R1, MAP2K2, HRAS, BAX | PTPN11, ABL1, NFKB1, RELA, SOS1, SOS2, GRB2, SHC4, AKT1, AKT3, PIK3CA, PIK3R2, MAPK3, BRAF |
| hsa04014 | Ras signaling pathway | 52,01720913 | 10 | 1,13112E-27 | 3,80279E-27 | MAP2K2, HRAS, PIK3CB, PIK3R1, SHC1, SHC2, IKBKB, RHOA, PLCG2 | MAPK3, GRB2, SOS1, SOS2, AKT1, AKT3, PIK3CA, PIK3R2, SHC4, NGF, NGFR, GAB1, PTPN11, RAC1, ABL1, NFKB1, RELA, MAPK8, MAPK9, MAPK10, CDC42, RAP1B, CALM3, PLCG1 |
| hsa04910 | Insulin signaling pathway | 73,72207407 | 10 | 1,80531E-27 | 1,2416E-24 | IKBKB, HRAS, MAP2K2, SHC1, SHC2, PIK3CB, PIK3R1, CRK, IRS1 | MAPK8, MAPK9, MAPK10, MAPK3, CALM3, SOS1, SOS2, SHC4, GRB2, BRAF, AKT1, AKT3, PIK3CA, PIK3R2, GSK3B, RAPGEF1, SH2B2, PDPK1 |
| hsa04917 | Prolactin signaling pathway | 109,986044 | 10 | 1,62382E-26 | 6,04594E-23 | PIK3CB, PIK3R1, MAP2K2, HRAS, SHC1, SHC2, FOXO3 | AKT1, AKT3, PIK3CA, PIK3R2, MAPK3, SHC4, GRB2, SOS1, SOS2, MAPK8, MAPK9, MAPK10, MAPK11, GSK3B, NFKB1, RELA |

| | | | | | | | |
|---|---|---|---|---|---|---|---|
| hsa05225 | Hepatocellular carcinoma | 60,62670565 | 10 | 5,93732E-26 | 2,12557E-23 | HRAS, MAP2K2, SHC1, SHC2, PIK3CB, PIK3R1, TP53, BAX, PLCG2 | BRAF, MAPK3, SHC4, GRB2, SOS1, SOS2, PIK3CA, PIK3R2, AKT1, AKT3, GSK3B, PLCG1, GAB1 |
| hsa05223 | Non-small cell lung cancer | 114,9854097 | 10 | 2,51323E-25 | 3,19405E-23 | FOXO3, PIK3CB, PIK3R1, HRAS, PLCG2, MAP2K2, TP53, BAX | AKT1, AKT3, PDPK1, PIK3CA, PIK3R2, PLCG1, SOS1, SOS2, GRB2, MAPK3, BRAF |
| hsa04072 | Phospholipase D signaling pathway | 71,67921641 | 10 | 8,046E-25 | 2,6278E-24 | MAP2K2, HRAS, PIK3CB, PIK3R1, SHC1, SHC2, RHOA, PLCG2 | MAPK3, GRB2, SOS1, SOS2, AKT1, AKT3, PIK3CA, PIK3R2, SHC4, GAB1, PTPN11, PLCG1 |
| hsa05211 | Renal cell carcinoma | 108,4148148 | 10 | 5,41992E-23 | 2,35491E-17 | CRK, PIK3CB, PIK3R1, JUN, MAP2K2, HRAS | PTPN11, RAC1, RAPGEF1, SOS1, SOS2, GRB2, GAB1, RAP1B, CDC42, AKT1, AKT3, PIK3CA, PIK3R2, MAPK3, BRAF |
| hsa04926 | Relaxin signaling pathway | 65,42273308 | 10 | 8,09645E-23 | 3,36419E-21 | NFKBIA, PIK3CB, PIK3R1, ATF4, HRAS, MAP2K2, SHC1, SHC2, JUN, MAP2K7 | AKT1, AKT3, NFKB1, RELA, MAPK3, PIK3CA, PIK3R2, SHC4, SOS1, SOS2, MAPK11, MAPK8, MAPK9, MAPK10, GRB2 |
| hsa04510 | Focal adhesion | 48,72575947 | 10 | 9,28674E-23 | 2,53741E-20 | PIK3CB, PIK3R1, RHOA, HRAS, SHC1, SHC2, CRK, JUN | CDC42, AKT1, AKT3, PDPK1, PIK3CA, PIK3R2, MAPK3, BRAF, SOS1, SOS2, GRB2, SHC4, RAPGEF1, RAP1B, MAPK8, MAPK9, MAPK10, RAC1, GSK3B |

| | | | | | | | |
|---|---|---|---|---|---|---|---|
| hsa05226 | Gastric cancer | 63,03186908 | 10 | 1,09564E-22 | 5,43048E-20 | HRAS, MAP2K2, SHC1, SHC2, PIK3CB, PIK3R1, TP53, BAX | BRAF, MAPK3, SHC4, GRB2, SOS1, SOS2, PIK3CA, PIK3R2, AKT1, AKT3, GSK3B, GAB1 |
| hsa04664 | Fc epsilon RI signaling pathway | 103,2522046 | 10 | 6,01692E-22 | 1,00366E-20 | MAP2K7, PIK3CB, PIK3R1, PLCG2, MAP2K2, HRAS | AKT1, AKT3, MAPK11, PDPK1, PIK3CA, PIK3R2, PLCG1, MAPK3, SOS1, SOS2, GRB2, MAPK8, MAPK9, MAPK10, RAC1 |
| hsa05210 | Colorectal cancer | 82,90544662 | 10 | 8,59585E-22 | 1,37313E-19 | PIK3CB, PIK3R1, RHOA, MAP2K2, JUN, TP53, BAX, HRAS | PIK3CA, PIK3R2, BRAF, MAPK8, MAPK9, MAPK10, RAC1, MAPK3, GSK3B, AKT1, AKT3, SOS1, SOS2, GRB2 |
| hsa05213 | Endometrial cancer | 114,1208577 | 10 | 1,57177E-21 | 1,91677E-19 | HRAS, FOXO3, PIK3CB, PIK3R1, MAP2K2, TP53, BAX | GSK3B, PDPK1, SOS1, SOS2, GRB2, AKT1, AKT3, PIK3CA, PIK3R2, MAPK3, BRAF |
| hsa04650 | Natural killer cell mediated cytotoxicity | 65,99162641 | 10 | 2,27911E-21 | 2,49875E-19 | PLCG2, PIK3CB, PIK3R1, MAP2K2, HRAS, SHC1, SHC2 | PLCG1, PTPN11, PIK3CA, PIK3R2, MAPK3, BRAF, SOS1, SOS2, GRB2, SHC4, RAC1 |
| hsa04068 | FoxO signaling pathway | 59,75619714 | 10 | 3,54221E-21 | 2,14896E-19 | IKBKB, FOXO3, HRAS, MAP2K2, PIK3CB, PIK3R1, IRS1 | MAPK8, MAPK9, MAPK10, MAPK3, SOS1, SOS2, BRAF, AKT1, AKT3, PIK3CA, PIK3R2, PDPK1, GRB2, MAPK11 |

| | | | | | | | |
|---|---|---|---|---|---|---|---|
| hsa05205 | Proteoglycans in cancer | 43,95195195 | 10 | 8,24803E-21 | 2,70634E-18 | RHOA, PIK3CB, PIK3R1, CAMK2A, CAMK2D, CAMK2G, HRAS, MAP2K2, TP53, PLCG2 | PIK3CA, PIK3R2, AKT1, AKT3, GAB1, RAC1, CDC42, BRAF, MAPK3, GRB2, PDPK1, FRS2, PTPN11, SOS1, SOS2, PLCG1, MAPK11 |
| hsa05224 | Breast cancer | 59,28935185 | 10 | 8,26464E-21 | 8,31491E-19 | HRAS, MAP2K2, SHC1, SHC2, JUN, PIK3CB, PIK3R1, TP53, BAX | BRAF, MAPK3, SHC4, GRB2, SOS1, SOS2, PIK3CA, PIK3R2, AKT1, AKT3, GSK3B |
| hsa05231 | Choline metabolism in cancer | 82,34036568 | 10 | 4,76474E-20 | 1,06833E-18 | HRAS, MAP2K2, PIK3CB, PIK3R1, JUN | MAPK8, MAPK9, MAPK10, MAPK3, SOS1, SOS2, AKT1, AKT3, PIK3CA, PIK3R2, PDPK1, GRB2, PLCG1, RAC1 |
| hsa04931 | Insulin resistance | 94,32438238 | 10 | 9,5119E-20 | 9,5119E-20 | IKBKB, IRS1, PIK3CB, PIK3R1, NFKBIA, RPS6KA1 | MAPK8, MAPK9, MAPK10, PIK3CA, PIK3R2, GSK3B, NFKB1, RELA, PTPN11, PDPK1, AKT1, AKT3 |
| hsa04625 | C-type lectin receptor signaling pathway | 115,409319 | 10 | 2,87919E-19 | 4,90687E-16 | HRAS, IKBKB, RHOA, MAPKAPK2, NFKBIA, PLCG2, PIK3CB, PIK3R1, JUN | NFKB1, RELA, PTPN11, MAPK3, PIK3CA, PIK3R2, AKT1, AKT3, CALM3, MAPK11, MAPK8, MAPK9, MAPK10 |
| hsa04668 | TNF signaling pathway | 103,2025641 | 10 | 2,96586E-19 | 4,92279E-14 | MAP2K7, IKBKB, NFKBIA, ATF4, JUN, PIK3CB, PIK3R1 | MAPK11, MAPK8, MAPK9, MAPK10, NFKB1, RELA, MAPK3, RPS6KA5, PIK3CA, PIK3R2, AKT1, AKT3 |

| | | | | | | | |
|---|---|---|---|---|---|---|---|
| hsa05167 | Kaposi's sarcoma-associated herpesvirus infection | 67,68092486 | 10 | 4,35529E-19 | 2,30737E-17 | IKBKB, NFKBIA, PIK3CB, PIK3R1, BAX, TP53, HRAS, MAP2K2, JUN, PLCG2, MAPKAPK2, MAP2K7 | MAPK8, MAPK9, MAPK10, AKT1, AKT3, PIK3CA, PIK3R2, NFKB1, RELA, GSK3B, PLCG1, MAPK3, RAC1, MAPK11, CALM3 |
| hsa05160 | Hepatitis C | 78,33719893 | 10 | 6,01622E-19 | 6,01622E-19 | NFKBIA, IKBKB, HRAS, PIK3CB, PIK3R1, TP53, MAP2K2, YWHAE, BAX | NFKB1, RELA, MAPK3, GRB2, SOS1, SOS2, BRAF, PIK3CA, PIK3R2, AKT1, AKT3, GSK3B |
| hsa04915 | Estrogen signaling pathway | 59,13535354 | 10 | 2,29351E-18 | 4,78807E-17 | JUN, HRAS, MAP2K2, ATF4, PIK3CB, PIK3R1, SHC1, SHC2 | MAPK3, PIK3CA, PIK3R2, AKT1, AKT3, SHC4, GRB2, SOS1, SOS2, CALM3 |
| hsa04660 | T cell receptor signaling pathway | 89,19445983 | 10 | 3,17563E-18 | 3,17563E-18 | NFKBIA, NFKBIB, IKBKB, PIK3CB, PIK3R1, HRAS, JUN, RHOA, MAP2K2, MAP2K7 | NFKBIE, NFKB1, RELA, AKT1, AKT3, PDPK1, GRB2, PIK3CA, PIK3R2, SOS1, SOS2, PLCG1, CDC42, MAPK3, MAPK9, MAPK11, GSK3B |
| hsa05169 | Epstein-Barr virus infection | 48,14473684 | 10 | 4,45148E-18 | 4,99677E-17 | TP53, NFKBIA, NFKBIB, MAP2K7, JUN, PIK3CB, PIK3R1, PLCG2, BAX, IRAK4, IKBKB | NFKBIE, NFKB1, RELA, MAPK8, MAPK9, MAPK10, MAPK11, PIK3CA, PIK3R2, AKT1, AKT3, RAC1 |
| hsa05200 | Pathways in cancer | 19,56530628 | 10 | 6,93927E-18 | 3,05842E-16 | CRK, PIK3CB, PIK3R1, IKBKB, HRAS, MAP2K2, NFKBIA, PLCG2, | ABL1, PIK3CA, PIK3R2, GSK3B, GRB2, SOS1, SOS2, BRAF, NFKB1, RELA, PLCG1, RAC1, MAPK8, MAPK9, |

| | | | | | | | |
|---|---|---|---|---|---|---|---|
| | | | | | | RHOA, JUN, TP53, BAX, CAMK2A, CAMK2D, CAMK2G | MAPK10, CDC42, MAPK3, AKT1, AKT3, RPS6KA5, CALM3 |
| hsa04062 | Chemokine signaling pathway | 45,75949976 | 10 | 3,12979E-17 | 6,48616E-15 | RHOA, PIK3CB, PIK3R1, SHC1, SHC2, CRK, HRAS, IKBKB, NFKBIA, NFKBIB, FOXO3 | PIK3CA, PIK3R2, SHC4, GRB2, CDC42, RAP1B, MAPK3, AKT1, AKT3, BRAF, RAC1, NFKB1, RELA, SOS1, SOS2, GSK3B |
| hsa04620 | Toll-like receptor signaling pathway | 106,057971 | 10 | 3,91907E-17 | 3,91863E-14 | IRAK4, IKBKB, NFKBIA, PIK3CB, PIK3R1, MAP2K7, MAP2K2, JUN | MAPK3, NFKB1, MAPK11, RAC1, MAPK8, MAPK9, MAPK10, AKT1, AKT3, PIK3CA, PIK3R2, RELA |
| hsa05215 | Prostate cancer | 64,81320451 | 10 | 5,27706E-17 | 2,84838E-16 | NFKBIA, HRAS, TP53, IKBKB, ATF4, PIK3CB, PIK3R1, MAP2K2 | NFKB1, RELA, PDPK1, SOS1, SOS2, GRB2, GSK3B, AKT1, AKT3, PIK3CA, PIK3R2, MAPK3, BRAF |
| hsa04010 | MAPK signaling pathway | 44,35151515 | 10 | 5,92863E-17 | 2,06716E-13 | IKBKB, MAP2K2, HRAS, MAP2K5, MAPKAPK2, ATF4, TP53, MAP2K7, CRK, JUN, MAP3K1, RPS6KA1, IRAK4 | NFKB1, RELA, MAPK3, BRAF, RAP1B, SOS1, SOS2, GRB2, NGFR, MAPK7, RPS6KA5, MAPK11, AKT1, AKT3, CDC42, RAC1, MAPK8, MAPK9, MAPK10, NGF |
| hsa05131 | Shigellosis | 104,3193158 | 10 | 9,53339E-17 | 6,09537E-12 | CRK, RIPK2, IKBKB, NFKBIA, NFKBIB | RAC1, ABL1, CDC42, NFKB1, RELA, MAPK8, MAPK9, MAPK10, MAPK3, MAPK11 |
| hsa05206 | MicroRNAs in cancer | 53,07018907 | 10 | 1,49143E-16 | 4,31076E-16 | CRK, HRAS, IRS1, RHOA, IKBKB, TP53, PLCG2, MAP2K2, SHC1 | ABL1, PIK3R2, MAPK7, PIK3CA, NFKB1, GRB2, RPS6KA5, PLCG1, SOS1, SOS2, SHC4 |
| hsa04380 | Osteoclast differentiation | 83,3960114 | 10 | 1,94843E-16 | 1,79108E-13 | IKBKB, NFKBIA, PIK3CB, PIK3R1, JUN, | NFKB1, RELA, MAPK8, MAPK9, MAPK10, |

| | | | | | | | |
|---|---|---|---|---|---|---|---|
| | | | | | | PLCG2, CAMK4, MAP2K7 | MAPK11, PIK3CA, PIK3R2, AKT1, AKT3, GRB2, MAPK3, RAC1 |
| hsa05161 | Hepatitis B | 50,72337744 | 10 | 2,75526E-16 | 5,19785E-13 | BAX, MAP2K2, ATF4, HRAS, MAP3K1, NFKBIA, JUN, TP53, PIK3CB, PIK3R1, IKBKB | MAPK3, NFKB1, RELA, GRB2, MAPK8, MAPK9, MAPK10, PIK3CA, PIK3R2, AKT1, AKT3 |
| hsa05142 | Chagas disease (American trypanosomiasis) | 92,43789474 | 10 | 3,72724E-16 | 3,41946E-14 | IRAK4, JUN, NFKBIA, IKBKB, PIK3CB, PIK3R1 | MAPK8, MAPK9, MAPK10, MAPK11, MAPK3, NFKB1, RELA, PIK3CA, PIK3R2, AKT1, AKT3 |
| hsa05120 | Epithelial cell signaling in Helicobacter pylori infection | 120,0902564 | 10 | 2,00628E-15 | 2,59249E-12 | JUN, IKBKB, NFKBIA, PLCG2 | MAPK11, MAPK8, MAPK9, MAPK10, NFKB1, RELA, CDC42, RAC1, PLCG1, PTPN11 |
| hsa04210 | Apoptosis | 59,25506073 | 10 | 2,54896E-15 | 6,43588E-15 | TP53, NFKBIA, IKBKB, PIK3CB, PIK3R1, BAX, JUN, ATF4, HRAS, MAP2K2 | NFKB1, RELA, AKT1, AKT3, PIK3CA, PIK3R2, NGF, MAPK8, MAPK9, MAPK10, PDPK1, MAPK3 |
| hsa05418 | Fluid shear stress and atherosclerosis | 67,0351145 | 10 | 2,54896E-15 | 9,87722E-12 | JUN, PIK3CB, PIK3R1, MAP2K7, MAP2K5, RHOA, TP53, IKBKB | MAPK11, AKT1, AKT3, MAPK8, MAPK9, MAPK10, PIK3CA, PIK3R2, MAPK7, RAC1, NFKB1, RELA, CALM3 |
| hsa05212 | Pancreatic cancer | 92,43789474 | 10 | 2,62162E-15 | 2,62162E-15 | PIK3CB, PIK3R1, IKBKB, TP53, BAX | PIK3CA, PIK3R2, AKT1, AKT3, RAC1, NFKB1, RELA, BRAF, MAPK3, MAPK8, MAPK9, MAPK10, CDC42 |

| | | | | | | | |
|---|---|---|---|---|---|---|---|
| hsa05164 | Influenza A | 61,36687631 | 10 | 3,79134E-15 | 4,40657E-14 | IKBKB, JUN, NFKBIA, NFKBIB, PIK3CB, PIK3R1, IRAK4, MAP2K7, MAP2K2 | NFKB1, RELA, PIK3CA, PIK3R2, AKT1, AKT3, MAPK11, MAPK8, MAPK9, MAPK10, GSK3B, MAPK3 |
| hsa04662 | B cell receptor signaling pathway | 141,410628 | 10 | 4,57686E-15 | 3,95783E-14 | NFKBIA, NFKBIB, IKBKB, PIK3CB, PIK3R1, JUN, HRAS, PLCG2, MAP2K2 | NFKBIE, NFKB1, RELA, GSK3B, AKT1, AKT3, PIK3CA, PIK3R2, RAC1, MAPK3, SOS1, SOS2, GRB2 |
| hsa04932 | Non-alcoholic fatty liver disease (NAFLD) | 57,91848041 | 10 | 1,26364E-14 | 1,26364E-14 | IRS1, PIK3CB, PIK3R1, BAX, ATF4, JUN, IKBKB | AKT1, AKT3, NFKB1, RELA, PIK3CA, PIK3R2, GSK3B, MAPK8, MAPK9, MAPK10, CDC42, RAC1 |
| hsa04150 | mTOR signaling pathway | 57,06042885 | 10 | 1,28548E-14 | 1,28548E-14 | IRS1, PIK3CB, PIK3R1, HRAS, MAP2K2, RHOA, RPS6KA1, IKBKB | PIK3CA, PIK3R2, BRAF, MAPK3, PDPK1, AKT1, AKT3, GRB2, SOS1, SOS2, GSK3B |
| hsa05221 | Acute myeloid leukemia | 76,22916667 | 10 | 1,36744E-14 | 1,61045E-12 | HRAS, IKBKB, PIK3CB, PIK3R1, MAP2K2 | NFKB1, RELA, SOS1, SOS2, GRB2, AKT1, AKT3, PIK3CA, PIK3R2, MAPK3, BRAF |
| hsa04933 | AGE-RAGE signaling pathway in diabetic complications | 127,2695652 | 10 | 1,55433E-14 | 1,91044E-12 | HRAS, JUN, BAX, PIK3CB, PIK3R1, PLCG2 | NFKB1, RELA, MAPK8, MAPK9, MAPK10, MAPK11, MAPK3, AKT1, AKT3, PIK3CA, PIK3R2, CDC42, RAC1, PLCG1 |

| | | | | | | | |
|---|---|---|---|---|---|---|---|
| hsa04658 | Th1 and Th2 cell differentiation | 114,7921569 | 10 | 1,72671E-14 | 3,68715E-10 | NFKBIA, NFKBIB, IKBKB, JUN | NFKBIE, NFKB1, RELA, MAPK3, MAPK11, MAPK8, MAPK9, MAPK10, PLCG1 |
| hsa04657 | IL-17 signaling pathway | 72,49532196 | 10 | 1,77766E-14 | 1,05407E-11 | IKBKB, NFKBIA, JUN | NFKB1, RELA, MAPK3, MAPK7, MAPK8, MAPK11, MAPK9, MAPK10, GSK3B |
| hsa04659 | Th17 cell differentiation | 97,57333333 | 10 | 5,07321E-14 | 4,98311E-10 | NFKBIA, NFKBIB, IKBKB, JUN | NFKBIE, NFKB1, RELA, MAPK3, MAPK11, MAPK8, MAPK9, MAPK10, PLCG1 |
| hsa04071 | Sphingolipid signaling pathway | 61,62526316 | 10 | 1,07874E-13 | 7,30978E-12 | MAP2K2, PIK3CB, PIK3R1, HRAS, RHOA, BAX, TP53 | MAPK3, AKT1, AKT3, PIK3CA, PIK3R2, PDPK1, MAPK8, MAPK9, MAPK10, MAPK11, NFKB1, RELA, RAC1 |
| hsa04370 | VEGF signaling pathway | 78,84713805 | 10 | 1,34036E-13 | 8,35396E-13 | PLCG2, MAPKAPK2, PIK3CB, PIK3R1, SHC2, MAP2K2, HRAS | CDC42, PLCG1, AKT1, AKT3, PIK3CA, PIK3R2, MAPK11, MAPK3, RAC1 |
| hsa04024 | cAMP signaling pathway | 48,14473684 | 10 | 1,52195E-13 | 1,52195E-13 | PIK3CB, PIK3R1, MAP2K2, RHOA, CAMK4, CAMK2A, CAMK2D, CAMK2G, NFKBIA, JUN | PIK3CA, PIK3R2, AKT1, AKT3, MAPK3, CALM3, RAP1B, BRAF, MAPK8, MAPK9, MAPK10, NFKB1, RELA, RAC1 |
| hsa04920 | Adipocytokine signaling pathway | 125,9010753 | 10 | 1,60425E-13 | 8,38041E-12 | NFKBIA, NFKBIB, IKBKB, IRS1 | NFKB1, RELA, NFKBIE, PTPN11, |

| | | | | | | | |
|---|---|---|---|---|---|---|---|
| | | | | | | | MAPK8, MAPK9, MAPK10, AKT1, AKT3 |
| hsa04151 | PI3K-Akt signaling pathway | 27,07180091 | 10 | 1,61506E-13 | 7,52106E-13 | PIK3CB, PIK3R1, IRS1, FOXO3, TP53, IKBKB, YWHAE, HRAS, MAP2K2, ATF4 | AKT1, AKT3, PDPK1, PIK3CA, PIK3R2, NGFR, GSK3B, NFKB1, RELA, NGF, GRB2, SOS1, SOS2, MAPK3, RAC1 |
| hsa04140 | Autophagy - animal | 56,36456996 | 10 | 1,867E-13 | 1,867E-13 | IRS1, PIK3CB, PIK3R1, HRAS, MAP2K2 | PIK3CA, PIK3R2, PDPK1, AKT1, AKT3, MAPK3, MAPK8, MAPK9, MAPK10 |
| hsa04622 | RIG-I-like receptor signaling pathway | 108,4148148 | 10 | 4,53341E-13 | 8,31561E-11 | IKBKB, NFKBIA, NFKBIB, MAP3K1 | NFKB1, RELA, MAPK8, MAPK9, MAPK10, MAPK11 |
| hsa05133 | Pertussis | 84,84637681 | 10 | 7,59242E-13 | 4,96116E-08 | RHOA, JUN, IRAK4 | CALM3, NFKB1, RELA, MAPK11, MAPK3, MAPK8, MAPK9, MAPK10 |
| hsa04912 | GnRH signaling pathway | 110,0451128 | 10 | 7,73612E-13 | 2,4581E-10 | ATF4, CAMK2A, CAMK2D, CAMK2G, MAP3K1, MAP2K2, HRAS, MAP2K7, JUN | CDC42, MAPK11, MAPK7, CALM3, MAPK3, SOS1, SOS2, GRB2, MAPK8, MAPK9, MAPK10 |
| hsa04930 | Type II diabetes mellitus | 122,5502392 | 10 | 7,93221E-13 | 7,93221E-13 | PIK3CB, PIK3R1, IRS1, IKBKB | PIK3CA, PIK3R2, MAPK3, MAPK8, MAPK9, MAPK10 |
| hsa05145 | Toxoplasmosis | 73,64025157 | 10 | 1,01422E-12 | 3,47907E-10 | IRAK4, IKBKB, NFKBIA, NFKBIB | NFKB1, RELA, MAPK3, MAPK8, MAPK9, MAPK10, MAPK11, PDPK1, AKT1, AKT3 |

| | | | | | | | |
|---|---|---|---|---|---|---|---|
| hsa05168 | Herpes simplex infection | 56,40077071 | 10 | 1,17419E-12 | 2,28225E-09 | JUN, NFKBIA, NFKBIB, TP53, IKBKB | MAPK8, MAPK9, MAPK10, NFKB1, RELA, PTPN11 |
| hsa05100 | Bacterial invasion of epithelial cells | 58,6026026 | 10 | 1,19393E-12 | 5,06176E-12 | CRK, PIK3CB, PIK3R1, SHC1, SHC2, RHOA | RAC1, CDC42, PIK3CA, PIK3R2, GAB1, SHC4 |
| hsa04621 | NOD-like receptor signaling pathway | 52,74234234 | 10 | 1,489E-12 | 2,20055E-09 | RIPK2, IKBKB, JUN, IRAK4, RHOA, NFKBIA, NFKBIB | MAPK8, MAPK9, MAPK10, MAPK3, MAPK11, NFKB1, RELA |
| hsa05132 | Salmonella infection | 74,10632911 | 10 | 1,72265E-12 | 8,66975E-08 | JUN | RAC1, CDC42, NFKB1, RELA, MAPK8, MAPK9, MAPK10, MAPK3, MAPK11 |
| hsa04914 | Progesterone-mediated oocyte maturation | 79,00674764 | 10 | 1,91044E-12 | 1,91044E-12 | PIK3CB, PIK3R1, RPS6KA1 | PIK3CA, PIK3R2, AKT1, AKT3, MAPK3, MAPK8, MAPK11, MAPK9, MAPK10, BRAF |
| hsa05203 | Viral carcinogenesis | 65,4122905 | 10 | 2,57253E-12 | 2,094E-09 | TP53, JUN, BAX, ATF4, PIK3CB, PIK3R1, HRAS, YWHAE, RHOA, NFKBIA, MAPKAPK2 | MAPK3, PIK3CA, PIK3R2, NFKB1, RELA, CDC42, RAC1, GRB2 |
| hsa05218 | Melanoma | 65,70594837 | 10 | 3,04822E-12 | 2,14277E-10 | TP53, PIK3CB, PIK3R1, MAP2K2, HRAS, BAX | BRAF, AKT1, AKT3, PIK3CA, PIK3R2, MAPK3 |
| hsa05166 | HTLV-I infection | 29,50145577 | 10 | 3,44856E-12 | 6,37825E-11 | HRAS, ATF4, NFKBIA, IKBKB, MAP3K1, JUN, PIK3CB, PIK3R1, BAX, MAP2K2, TP53 | NFKB1, RELA, MAPK8, MAPK9, MAPK10, PIK3CA, PIK3R2, AKT1, AKT3, MAPK3 |
| hsa04213 | Longevity regulating pathway - multiple species | 94,60018467 | 10 | 6,50193E-12 | 6,50193E-12 | FOXO3, PIK3CB, PIK3R1, IRS1, HRAS | AKT1, AKT3, PIK3CA, PIK3R2 |

| | | | | | | | |
|---|---|---|---|---|---|---|---|
| hsa04919 | Thyroid hormone signaling pathway | 56,53693868 | 10 | 6,50817E-12 | 6,50817E-12 | PIK3CB, PIK3R1, PLCG2, HRAS, MAP2K2, TP53 | AKT1, AKT3, PDPK1, PIK3CA, PIK3R2, GSK3B, PLCG1, MAPK3 |
| hsa04630 | Jak-STAT signaling pathway | 34,60047281 | 10 | 7,36204E-12 | 5,71421E-11 | PIK3CB, PIK3R1, HRAS | AKT1, AKT3, PIK3CA, PIK3R2, SOS1, SOS2, GRB2, PTPN11 |
| hsa04152 | AMPK signaling pathway | 55,5182551 | 10 | 8,19451E-12 | 8,19451E-12 | PIK3CB, PIK3R1, IRS1, FOXO3 | AKT1, AKT3, PIK3CA, PIK3R2, PDPK1 |
| hsa04066 | HIF-1 signaling pathway | 47,13687601 | 10 | 8,41733E-12 | 5,28596E-11 | PIK3CB, PIK3R1, PLCG2, CAMK2A, CAMK2D, CAMK2G, MAP2K2 | PIK3CA, PIK3R2, AKT1, AKT3, PLCG1, NFKB1, RELA, MAPK3 |
| hsa04015 | Rap1 signaling pathway | 30,11522634 | 10 | 8,94991E-12 | 1,96269E-09 | MAP2K2, CRK, RHOA, PIK3CB, PIK3R1, HRAS | MAPK3, RAP1B, NGF, NGFR, RAC1, MAPK11, BRAF, RAPGEF1, CDC42, PIK3CA, PIK3R2, AKT1, AKT3, CALM3, PLCG1 |
| hsa05162 | Measles | 61,95132275 | 10 | 1,16101E-11 | 4,94073E-10 | IRAK4, NFKBIA, NFKBIB, PIK3CB, PIK3R1, TP53 | NFKB1, RELA, PIK3CA, PIK3R2, AKT1, AKT3, TP73, GSK3B |
| hsa04211 | Longevity regulating pathway | 65,75866496 | 10 | 1,28478E-11 | 7,04412E-11 | FOXO3, PIK3CB, PIK3R1, IRS1, HRAS, ATF4, TP53, BAX, CAMK4 | AKT1, AKT3, PIK3CA, PIK3R2, NFKB1, RELA |
| hsa04137 | Mitophagy - animal | 73,02352103 | 10 | 1,90543E-11 | 9,51744E-11 | ATF4, TP53, HRAS, JUN, FOXO3 | RELA, MAPK8, MAPK9, MAPK10 |
| hsa05230 | Central carbon metabolism in cancer | 63,24197531 | 10 | 2,86655E-11 | 1,37328E-10 | HRAS, MAP2K2, PIK3CB, PIK3R1, TP53 | NTRK3, MAPK3, PIK3CA, PIK3R2, AKT1, AKT3 |
| hsa05222 | Small cell lung cancer | 77,61515152 | 10 | 2,92478E-11 | 6,5469E-11 | PIK3CB, PIK3R1, IKBKB, NFKBIA, BAX, TP53 | AKT1, AKT3, PIK3CA, PIK3R2, NFKB1, RELA |
| hsa05219 | Bladder cancer | 178,4878049 | 10 | 4,43836E-11 | 4,80102E-07 | HRAS, TP53, MAP2K2 | RPS6KA5, MAPK3, BRAF |

| | | | | | | | |
|---|---|---|---|---|---|---|---|
| hsa05165 | Human papillomavirus infection | 19,2224849 | 10 | 6,88041E-11 | 7,40725E-10 | PIK3CB, PIK3R1, TP53, IKBKB, HRAS, MAP2K2, BAX | AKT1, AKT3, PIK3CA, PIK3R2, GSK3B, NFKB1, RELA, GRB2, SOS1, SOS2, MAPK3, CDC42 |
| hsa04750 | Inflammatory mediator regulation of TRP channels | 69,13090418 | 10 | 1,23453E-10 | 1,23453E-10 | CAMK2A, CAMK2D, CAMK2G, PLCG2, PIK3CB, PIK3R1 | NGF, MAPK8, MAPK11, MAPK9, MAPK10, PLCG1, PIK3CA, PIK3R2, CALM3 |
| hsa04923 | Regulation of lipolysis in adipocytes | 100,4759725 | 10 | 2,77216E-10 | 2,77216E-10 | PIK3CB, PIK3R1, IRS1 | PIK3CA, PIK3R2, AKT1, AKT3 |
| hsa04810 | Regulation of actin cytoskeleton | 26,22939068 | 10 | 3,034E-10 | 6,49345E-08 | RHOA, MAP2K2, PIK3CB, PIK3R1, CRK, HRAS | CDC42, RAC1, MAPK3, BRAF, PIK3CA, PIK3R2, SOS1, SOS2 |
| hsa05216 | Thyroid cancer | 203,2777778 | 10 | 4,44267E-10 | 1,27677E-05 | HRAS, MAP2K2, TP53, BAX | BRAF, MAPK3 |
| hsa04623 | Cytosolic DNA-sensing pathway | 108,4148148 | 10 | 4,82397E-10 | 3,14794E-08 | IKBKB, NFKBIA, NFKBIB | NFKB1, RELA |
| hsa04218 | Cellular senescence | 44,93508772 | 10 | 5,59748E-10 | 3,67931E-09 | TP53, PIK3CB, PIK3R1, FOXO3, HRAS, MAP2K2, MAPKAPK2 | PIK3CA, PIK3R2, AKT1, AKT3, MAPK3, MAPK11, NFKB1, RELA, CALM3 |
| hsa05140 | Leishmaniasis | 84,84637681 | 10 | 6,53802E-10 | 4,43849E-08 | NFKBIA, NFKBIB, IRAK4, JUN | NFKB1, RELA, MAPK3, MAPK11 |
| hsa04666 | Fc gamma R-mediated phagocytosis | 45,17283951 | 10 | 9,49979E-10 | 6,00668E-09 | PIK3CB, PIK3R1, PLCG2, CRK | PIK3CA, PIK3R2, PLCG1, MAPK3, AKT1, AKT3, RAC1, CDC42 |
| hsa04310 | Wnt signaling pathway | 59,01612903 | 10 | 1,05448E-09 | 1,0873E-08 | CAMK2A, CAMK2D, CAMK2G, RHOA, JUN, TP53 | MAPK8, MAPK9, MAPK10, RAC1, GSK3B |
| hsa04530 | Tight junction | 49,44594595 | 10 | 1,84034E-09 | 4,26059E-08 | RHOA, MAP2K7, MAP3K1, JUN | CDC42, RAC1, MAPK8, MAPK9, MAPK10 |
| hsa04730 | Long-term depression | 138,0754717 | 10 | 2,57594E-09 | 1,97262E-05 | MAP2K2, HRAS | MAPK3, BRAF |

| | | | | | | | |
|---|---|---|---|---|---|---|---|
| hsa04720 | Long-term potentiation | 63,63478261 | 10 | 3,80173E-09 | 1,30425E-08 | CAMK4, CAMK2A, CAMK2D, CAMK2G, RPS6KA1, ATF4, MAP2K2, HRAS | CALM3, MAPK3, BRAF, RAP1B |
| hsa04960 | Aldosterone-regulated sodium reabsorption | 113,2817337 | 10 | 4,33551E-09 | 4,33551E-09 | IRS1, PIK3CB, PIK3R1 | PIK3CA, PIK3R2, PDPK1, MAPK3 |
| hsa04215 | Apoptosis - multiple species | 99,42934783 | 10 | 7,89756E-09 | 1,06891E-05 | BAX | MAPK8, MAPK9, MAPK10, NGFR |
| hsa04973 | Carbohydrate digestion and absorption | 132,8130672 | 10 | 1,04208E-08 | 1,04208E-08 | PIK3CB, PIK3R1 | PIK3CA, PIK3R2, AKT1, AKT3 |
| hsa04540 | Gap junction | 35,1996152 | 10 | 1,34123E-08 | 2,05752E-06 | MAP2K5, MAP2K2, HRAS | MAPK7, MAPK3, SOS1, SOS2, GRB2 |
| hsa05152 | Tuberculosis | 36,59 | 10 | 1,89658E-08 | 1,85272E-06 | CAMK2A, CAMK2D, CAMK2G, IRAK4, BAX, RHOA, RIPK2 | CALM3, MAPK3, NFKB1, RELA, AKT1, AKT3, MAPK8, MAPK9, MAPK10, MAPK11, IRAK2 |
| hsa04725 | Cholinergic synapse | 48,14473684 | 10 | 2,02929E-08 | 2,02929E-08 | ATF4, CAMK4, CAMK2A, CAMK2D, CAMK2G, PIK3CB, PIK3R1, HRAS | MAPK3, PIK3CA, PIK3R2, AKT1, AKT3 |
| hsa04360 | Axon guidance | 26,53509454 | 10 | 2,16256E-08 | 1,02481E-07 | RHOA, HRAS, CAMK2A, CAMK2D, CAMK2G, PIK3CB, PIK3R1, PLCG2 | MAPK3, RAC1, CDC42, ABL1, PIK3CA, PIK3R2, GSK3B, PLCG1, PTPN11 |
| hsa04921 | Oxytocin signaling pathway | 36,8135106 | 10 | 3,35515E-08 | 7,07596E-06 | HRAS, MAP2K2, CAMK4, CAMK2A, CAMK2D, CAMK2G, RHOA, MAP2K5, JUN | CALM3, MAPK3, MAPK7 |

| | | | | | | | |
|---|---|---|---|---|---|---|---|
| hsa04728 | Dopaminergic synapse | 38,19747716 | 10 | 5,14989E-08 | 5,14989E-08 | ATF4, CAMK2A, CAMK2D, CAMK2G | AKT1, AKT3, GSK3B, MAPK8, MAPK11, MAPK9, MAPK10, CALM3 |
| hsa04670 | Leukocyte transendothelial migration | 33,18820862 | 10 | 5,24911E-08 | 1,93793E-07 | PIK3CB, PIK3R1, RHOA, PLCG2 | PTPN11, PIK3CA, PIK3R2, MAPK11, RAC1, RAP1B, PLCG1, CDC42 |
| hsa04550 | Signaling pathways regulating pluripotency of stem cells | 38,51578947 | 10 | 7,25068E-08 | 7,25068E-08 | PIK3CB, PIK3R1, HRAS, MAP2K2 | GSK3B, PIK3CA, PIK3R2, AKT1, AKT3, GRB2, MAPK3, MAPK11 |
| hsa05034 | Alcoholism | 27,90087146 | 10 | 9,31807E-08 | 2,35278E-06 | ATF4, CAMK4, HRAS, SHC1, SHC2 | CALM3, BRAF, MAPK3, SHC4, GRB2, SOS1, SOS2 |
| hsa04371 | Apelin signaling pathway | 45,7375 | 10 | 1,01385E-07 | 0,001191624 | MAP2K2, HRAS, CAMK4 | AKT1, AKT3, MAPK3, CALM3 |
| hsa05321 | Inflammatory bowel disease (IBD) | 102,7087719 | 10 | 2,10959E-07 | 7,81113E-05 | JUN | NFKB1, RELA |
| hsa04064 | NF-kappa B signaling pathway | 54,20740741 | 10 | 2,35681E-07 | 1,32045E-05 | NFKBIA, IRAK4, IKBKB, PLCG2 | NFKB1, RELA, PLCG1 |
| hsa05030 | Cocaine addiction | 95,45217391 | 10 | 4,97715E-07 | 2,01192E-05 | JUN, ATF4 | NFKB1, RELA |
| hsa05031 | Amphetamine addiction | 52,15965788 | 10 | 5,20926E-07 | 4,68146E-05 | CAMK4, CAMK2A, CAMK2D, CAMK2G, ATF4, JUN | CALM3 |
| hsa04070 | Phosphatidylinositol signaling system | 46,97047497 | 10 | 5,50046E-07 | 5,50046E-07 | PLCG2, PIK3CB, PIK3R1 | PLCG1, CALM3, PIK3CA, PIK3R2 |
| hsa04261 | Adrenergic signaling in cardiomyocytes | 31,8173913 | 10 | 5,73063E-07 | 0,000231754 | ATF4, CAMK2A, CAMK2D, CAMK2G | AKT1, AKT3, MAPK3, CALM3, MAPK11, RPS6KA5 |
| hsa05146 | Amoebiasis | 68,87529412 | 10 | 8,09332E-07 | 6,09204E-06 | PIK3CB, PIK3R1 | NFKB1, RELA, PIK3CA, PIK3R2 |
| hsa04723 | Retrograde endocannabinoid signaling | 33,64597701 | 10 | 1,12605E-06 | 0,000545003 | | MAPK3, MAPK8, MAPK11, MAPK9, MAPK10 |
| hsa04922 | Glucagon signaling pathway | 36,15612648 | 10 | 1,27804E-06 | 0,000125403 | ATF4, CAMK2A, CAMK2D, CAMK2G | AKT1, AKT3, CALM3 |

| | | | | | | | |
|---|---|---|---|---|---|---|---|
| hsa04611 | Platelet activation | 35,99606493 | 10 | 1,57114E-06 | 1,57114E-06 | PLCG2, PIK3CB, PIK3R1, RHOA | RAP1B, PIK3CA, PIK3R2, AKT1, AKT3, MAPK11, MAPK3 |
| hsa04916 | Melanogenesis | 31,424584 | 10 | 1,70645E-06 | 7,90659E-05 | CAMK2A, CAMK2D, CAMK2G, MAP2K2, HRAS | CALM3, GSK3B, MAPK3 |
| hsa04934 | Cushing's syndrome | 25,65918654 | 10 | 1,78738E-06 | 1,87697E-05 | ATF4, CAMK2A, CAMK2D, CAMK2G, MAP2K2 | GSK3B, MAPK3, RAP1B, BRAF |
| hsa04726 | Serotonergic synapse | 57,171875 | 10 | 2,43866E-06 | 0,006186365 | HRAS | MAPK3, BRAF |
| hsa04270 | Vascular smooth muscle contraction | 61,66853933 | 10 | 3,20422E-06 | 0,000173695 | RHOA, MAP2K2 | CALM3, BRAF, MAPK3 |
| hsa05020 | Prion diseases | 114,34375 | 10 | 5,54019E-06 | 0,001155479 | BAX, MAP2K2 | MAPK3 |
| hsa00562 | Inositol phosphate metabolism | 50,51251079 | 10 | 7,47618E-06 | 7,47618E-06 | PLCG2, PIK3CB | PLCG1, PIK3CA |
| hsa04141 | Protein processing in endoplasmic reticulum | 17,6763285 | 10 | 1,42641E-05 | 0,001987583 | ATF4, MAP2K7, BAX | MAPK8, MAPK9, MAPK10 |
| hsa04022 | cGMP-PKG signaling pathway | 33,01353383 | 10 | 1,42846E-05 | 0,000778012 | RHOA, MAP2K2, ATF4, IRS1 | MAPK3, AKT1, AKT3, CALM3 |
| hsa05202 | Transcriptional misregulation in cancer | 15,42661397 | 10 | 1,46476E-05 | 0,000608699 | TP53, BAX | NGFR, NFKB1, RELA |
| hsa05014 | Amyotrophic lateral sclerosis (ALS) | 58,544 | 10 | 1,68298E-05 | 0,000108274 | BAX, TP53 | MAPK11, RAC1 |
| hsa05134 | Legionellosis | 73,91919192 | 10 | 1,88339E-05 | 6,30598E-05 | NFKBIA | NFKB1, RELA |
| hsa04520 | Adherens junction | 60,47933884 | 10 | 2,77168E-05 | 0,006679214 | RHOA | MAPK3, RAC1, CDC42 |
| hsa04110 | Cell cycle | 20,86386315 | 10 | 3,404E-05 | 0,000474639 | YWHAE, TP53 | GSK3B, ABL1 |
| hsa04114 | Oocyte meiosis | 25,71102328 | 10 | 4,05766E-05 | 0,000209923 | RPS6KA1, CAMK2A, CAMK2D, CAMK2G, YWHAE | MAPK3, CALM3 |
| hsa04713 | Circadian entrainment | 31,8173913 | 10 | 6,65957E-05 | 0,004164564 | CAMK2A, CAMK2D, CAMK2G | CALM3, MAPK3, RPS6KA5 |
| hsa04925 | Aldosterone synthesis and secretion | 33,49199085 | 10 | 6,65957E-05 | 0,004164564 | ATF4, CAMK4, CAMK2A, CAMK2D, CAMK2G | CALM3 |
| hsa04390 | Hippo signaling pathway | 18,0691358 | 10 | 8,01898E-05 | 0,004639423 | YWHAE | TP73, GSK3B |
| hsa04928 | Parathyroid hormone synthesis, secretion and action | 40,65555556 | 10 | 0,000136943 | 0,005935496 | RHOA, ATF4 | BRAF, MAPK3 |
| hsa04217 | Necroptosis | 18,0524206 | 10 | 0,000153536 | 0,001097604 | BAX, CAMK2A, CAMK2D, CAMK2G | MAPK8, MAPK9, MAPK10 |

| | | | | | | | |
|---|---|---|---|---|---|---|---|
| hsa05130 | Pathogenic Escherichia coli infection | 58,07936508 | 10 | 0,00015932 | 0,000200269 | RHOA | CDC42, ABL1 |
| hsa04115 | p53 signaling pathway | 29,36989967 | 10 | 0,000192385 | 0,00052814 | BAX, TP53 | TP73 |
| hsa04714 | Thermogenesis | 14,67121091 | 10 | 0,00022664 | 0,003153946 | RPS6KA1, HRAS | MAPK11, FRS2, GRB2, SOS1, SOS2 |
| hsa05217 | Basal cell carcinoma | 37,43222506 | 10 | 0,000367623 | 0,004103374 | TP53, BAX | GSK3B |
| hsa04971 | Gastric acid secretion | 35,352657 | 10 | 0,000495548 | 0,006683375 | CAMK2A, CAMK2D, CAMK2G | CALM3 |
| hsa04020 | Calcium signaling pathway | 17,6763285 | 10 | 0,000670534 | 0,005314531 | CAMK4, CAMK2A, CAMK2D, CAMK2G, PLCG2 | PLCG1, CALM3 |
| hsa04911 | Insulin secretion | 28,49318624 | 10 | 0,000709588 | 0,010421479 | ATF4, CAMK2A, CAMK2D, CAMK2G | |
| hsa04144 | Endocytosis | 14,19133807 | 10 | 0,000760451 | 0,006243417 | HRAS, RHOA | CDC42 |
| hsa05010 | Alzheimer's disease | 19,27480246 | 10 | 0,001176332 | 0,004372411 | PSEN2 | MAPK3, CALM3, GSK3B |
| hsa05110 | Vibrio cholerae infection | 25,81305115 | 10 | 0,002009405 | 0,011660354 | PLCG2 | PLCG1 |
| hsa04340 | Hedgehog signaling pathway | 56,29230769 | 2 | 0,002215854 | 0,002306028 | | GSK3B |
| hsa04962 | Vasopressin-regulated water reabsorption | 91,475 | 10 | 0,00250219 | 0,004874685 | ARHGDIG | ARHGDIA |
| hsa04350 | TGF-beta signaling pathway | 57,28375734 | 6 | 0,00334449 | 0,021606948 | RHOA | MAPK3 |
| hsa04972 | Pancreatic secretion | 36,45330012 | 10 | 0,003394377 | 0,012487115 | RHOA | RAP1B, RAC1 |
| hsa05416 | Viral myocarditis | 78,90026954 | 5 | 0,00340777 | 0,00913641 | | ABL1, RAC1 |
| hsa05323 | Rheumatoid arthritis | 50,99651568 | 2 | 0,004814402 | 0,004814402 | JUN | |
| hsa04120 | Ubiquitin mediated proteolysis | 18,34085213 | 10 | 0,007052993 | 0,025139894 | MAP3K1 | |
| hsa05016 | Huntington's disease | 17,6231186 | 10 | 0,009404782 | 0,03118457 | TP53, BAX | |
| hsa04330 | Notch signaling pathway | 221,7575758 | 2 | 0,010058306 | 0,010058306 | PSEN2 | |
| hsa04724 | Glutamatergic synapse | 23,49277689 | 1 | 0,015492544 | 0,015492544 | | MAPK3 |
| hsa04740 | Olfactory transduction | 64,19298246 | 1 | 0,044741685 | 0,044741685 | CAMK2A, CAMK2D, CAMK2G | CALM3 |
| hsa05310 | Asthma | 159,0869565 | 1 | 0,049895404 | 0,049895404 | | |

**Supplementary Table S2.** Enrichment analysis (KEGG pathways) of *BMI1*-correlated genes.

| ID | Pathway | Fold_Enrichment | occurrence | lowest_p | highest_p | Up_regulated | Down_regulated |
|---|---|---|---|---|---|---|---|
| hsa04010 | MAPK signaling pathway | 53,02898551 | 10 | 6,82809E-78 | 1,27341E-65 | RASGRF1, DUSP4, DUSP5, DUSP3, PTPRR, PTPN5, PPM1B, MAP4K4, STMN1, MAPT, CHUK, IKBKG, MAPK1, RAF1, BRAF, RASA1, NF1, RAP1A, PRKACA, PRKACB, RAPGEF2, RASA2, RRAS2, MRAS, RASGRP2, SOS1, SOS2, GNA12, GRB2, NTRK2, CACNA1A, CACNA1C, CACNA1E, CACNA1F, CACNA2D1, CACNB1, CACNB3, CACNG1, CACNA1H, CACNA2D2, CACNG3, CACNG2, NLK, RPS6KA5, ELK4, ATF2, MAPK14, MAPK11, MAPK13, PPM1A, AKT3, TAOK2, TAOK3, TAOK1, MAP3K7, MAP3K13, MAP4K2, STK4, TAB2, TAB1, GADD45G, TRAF6, CASP3, TGFB3, CDC42, RAC1, PAK1, PAK2, MAP3K2, MAPK8IP1, MAPK8, MAPK9, ARRB1, HSPA1L, HSPA8, PPP3CA, PPP3CB, PPP3R1, PPP3R2, MAP4K3, MKNK1, FGF9, NTF3, FGF16 | RASGRF2, DUSP1, DUSP2, DUSP6, DUSP7, DUSP8, DUSP9, DUSP10, DUSP16, PTPN7, PLA2G4A, NFKB1, NFKB2, RELA, RELB, IKBKB, MAP2K2, PRKCA, PRKCG, HRAS, RRAS, RASGRP3, RASGRP4, EGFR, FGFR3, FGFR2, FGFR4, NTRK1, PDGFRA, PDGFRB, CACNA1B, CACNA1S, CACNB2, CACNA1I, CACNG5, CACNG4, NR4A1, CDC25B, RPS6KA4, MAPKAPK3, MAPKAPK2, ATF4, MEF2C, DDIT3, TP53, ELK1, MAP2K6, MAP2K3, AKT2, MAP3K4, HSPB1, MAP3K6, MAP3K12, STK3, ECSIT, GADD45A, GADD45B, DAXX, CD14, FAS, IL1R1, TNFRSF1A, TGFB2, IL1B, RAC2, MAPK8IP2, MAP3K11, MAP2K7, MAPK10, ARRB2, HSPA2, HSPA6, JUND, JUN, NFATC3, NFATC1, FLNA, FLNB, FLNC, MAP3K1, MAP3K8, MAP4K1, FOS, SRF, MYC, RPS6KA1, RPS6KA2, RPS6KA3, MKNK2, BDNF, FGF1, FGF3, FGF5, FGF7, FGF8, FGF10, PDGFB, FGF18, FGF17, FGF19, FGF20 |

| | | | | | | | |
|---|---|---|---|---|---|---|---|
| hsa05161 | Hepatitis B | 67,87710145 | 10 | 9,90005E-25 | 7,17929E-23 | CASP3, RAF1, MAPK1, ATF2, GRB2, MAPK8, MAPK9, AKT3, CHUK, IKBKG, TGFB3 | PRKCA, PRKCG, MAP2K2, FOS, NFKB1, RELA, MYC, ELK1, ATF4, HRAS, MAP3K1, MAPK10, JUN, NFATC1, NFATC3, TP53, AKT2, IKBKB, FAS, TGFB2 |
| hsa05169 | Epstein-Barr virus infection | 34,64962121 | 10 | 4,1363E-24 | 7,59837E-22 | CHUK, TRAF6, TAB1, TAB2, MAP3K7, MAPK8, MAPK9, MAPK14, MAPK11, MAPK13, AKT3, GADD45G, RAC1, IKBKG, CASP3 | MYC, TP53, NFKB1, RELA, NFKB2, RELB, MAP2K7, MAP2K3, MAP2K6, MAPK10, JUN, AKT2, GADD45A, GADD45B, IKBKB, FAS |
| hsa04912 | GnRH signaling pathway | 79,29721362 | 10 | 3,58056E-22 | 3,40562E-20 | CDC42, MAPK14, MAPK11, MAPK13, MAP3K2, PRKACA, PRKACB, CACNA1C, CACNA1F, MAPK1, RAF1, SOS1, SOS2, GRB2, MAPK8, MAPK9 | PRKCA, ATF4, MAP2K3, MAP2K6, MAP3K1, MAP3K4, PLA2G4A, CACNA1S, ELK1, MAP2K2, HRAS, EGFR, MAP2K7, MAPK10, JUN |
| hsa05205 | Proteoglycans in cancer | 45,80256046 | 10 | 6,83522E-21 | 4,27626E-13 | AKT3, RAC1, CDC42, PAK1, MAPK1, RRAS2, MRAS, BRAF, RAF1, GRB2, SOS1, SOS2, PRKACA, PRKACB, MAPK14, MAPK11, MAPK13, CASP3 | AKT2, FLNA, FLNB, FLNC, ELK1, HRAS, RRAS, MAP2K2, TP53, PRKCA, MYC, FAS, TGFB2, PRKCG, EGFR |

| | | | | | | | |
|---|---|---|---|---|---|---|---|
| hsa04668 | TNF signaling pathway | 73,18 | 10 | 1,04391E-20 | 5,54129E-17 | MAP3K7, MAPK14, MAPK11, MAPK13, MAPK8, MAPK9, IKBKG, MAPK1, RPS6KA5, ATF2, CASP3, AKT3, CHUK, TAB1, TAB2 | TNFRSF1A, MAP2K7, MAP2K3, MAP2K6, MAPK10, IKBKB, NFKB1, RELA, MAP3K8, RPS6KA4, ATF4, JUN, FAS, IL1B, FOS, AKT2 |
| hsa04014 | Ras signaling pathway | 39,42356902 | 10 | 3,87046E-20 | 1,79906E-19 | MAPK1, RAF1, RRAS2, MRAS, GRB2, SOS1, SOS2, AKT3, FGF9, NTF3, FGF16, NTRK2, RAC1, CHUK, IKBKG, MAPK8, MAPK9, CDC42, RASGRP2, RASGRF1, RASA1, RASA2, NF1, STK4, RAP1A, PAK1, PAK2, PRKACA, PRKACB | MAP2K2, HRAS, RRAS, AKT2, BDNF, FGF1, FGF3, FGF5, FGF7, FGF8, FGF10, PDGFB, FGF18, FGF17, FGF19, FGF20, EGFR, FGFR3, FGFR2, FGFR4, NTRK1, PDGFRA, PDGFRB, RAC2, PRKCA, PRKCG, IKBKB, NFKB1, RELA, MAPK10, PLA2G4A, ELK1, RASGRP3, RASGRP4, RASGRF2 |
| hsa04722 | Neurotrophin signaling pathway | 49,37921727 | 10 | 6,14566E-20 | 2,16588E-17 | NTRK2, NTF3, GRB2, AKT3, BRAF, RAF1, SOS1, SOS2, MAPK1, RAP1A, RPS6KA5, MAPK8, MAPK9, MAPK14, MAPK11, MAPK13, CDC42, RAC1, TRAF6 | NTRK1, BDNF, AKT2, HRAS, MAP2K2, RPS6KA1, RPS6KA2, RPS6KA3, NFKB1, RELA, ATF4, MAPK10, TP53, MAP3K1, JUN, MAPKAPK2, MAP2K7, IKBKB |
| hsa04380 | Osteoclast differentiation | 65,04888889 | 10 | 7,14425E-20 | 1,0268E-17 | TRAF6, CHUK, IKBKG, MAP3K7, TAB1, TAB2, MAPK8, MAPK9, MAPK14, MAPK11, MAPK13, AKT3, PPP3CA, PPP3CB, PPP3R1, PPP3R2, GRB2, MAPK1, RAC1 | IKBKB, NFKB1, RELA, MAP2K6, MAPK10, NFATC1, AKT2, IL1B, IL1R1, FOS, JUN, JUND, TNFRSF1A, TGFB2, NFKB2, RELB, MAP2K7 |
| hsa04625 | C-type lectin receptor signaling pathway | 62,95053763 | 10 | 2,96484E-19 | 4,27744E-17 | RAF1, PAK1, RRAS2, MRAS, IKBKG, CHUK, MAPK1, PPP3CA, PPP3CB, PPP3R1, PPP3R2, AKT3, MAPK14, MAPK11, MAPK13, | NFKB1, HRAS, RRAS, RELA, IKBKB, IL1B, MAPKAPK2, NFATC1, NFATC3, AKT2, NFKB2, RELB, MAPK10, JUN |

| | | | | | | | |
|---|---|---|---|---|---|---|---|
| | | | | | | MAPK8, MAPK9 | |
| hsa04620 | Toll-like receptor signaling pathway | 76,36173913 | 10 | 6,96493E-19 | 5,71318E-17 | CHUK, TRAF6, MAPK1, MAP3K7, MAPK14, MAPK11, MAPK13, RAC1, MAPK8, MAPK9, TAB1, AKT3, TAB2, IKBKG | MAP3K8, NFKB1, IKBKB, MAPK10, IL1B, AKT2, CD14, MAP2K7, RELA, MAP2K3, MAP2K6, MAP2K2, FOS, JUN |
| hsa05145 | Toxoplasmosis | 59,8327044 | 10 | 1,0892E-18 | 1,9422E-18 | TRAF6, MAP3K7, TAB1, TAB2, CHUK, IKBKG, MAPK1, MAPK8, MAPK9, MAPK14, MAPK11, MAPK13, CASP3, AKT3, HSPA1L, HSPA8, TGFB3 | IKBKB, NFKB1, RELA, MAPK10, MAP2K3, MAP2K6, AKT2, HSPA2, HSPA6, TNFRSF1A, TGFB2 |
| hsa05160 | Hepatitis C | 49,61355932 | 10 | 1,93407E-18 | 3,80597E-16 | CHUK, IKBKG, TRAF6, MAPK1, GRB2, SOS1, SOS2, BRAF, RAF1, AKT3, CASP3 | NFKB1, RELA, IKBKB, EGFR, HRAS, AKT2, TP53, TNFRSF1A, MAP2K2, FAS, MYC |
| hsa05200 | Pathways in cancer | 14,83092067 | 10 | 2,19799E-18 | 1,39671E-15 | CASP3, CHUK, IKBKG, GRB2, SOS1, SOS2, BRAF, RAF1, STK4, RAC1, MAPK8, MAPK9, CDC42, TGFB3, TRAF6, MAPK1, AKT3, FGF9, FGF16, PRKACA, PRKACB, GNA12, RASGRP2, GADD45G, RPS6KA5 | EGFR, PDGFRA, PDGFRB, FGFR3, FGFR2, FGFR4, NTRK1, PDGFB, IKBKB, HRAS, MAP2K2, NFKB1, NFKB2, RELA, RAC2, MAPK10, JUN, FOS, MYC, TP53, PRKCA, PRKCG, TGFB2, FAS, AKT2, FGF1, FGF3, FGF5, FGF7, FGF8, FGF10, FGF18, FGF17, FGF19, FGF20, RASGRP3, RASGRP4, GADD45A, GADD45B, ELK1 |
| hsa05167 | Kaposi's sarcoma-associated herpesvirus infection | 39,4805395 | 10 | 7,2365E-18 | 1,69746E-17 | CHUK, IKBKG, MAPK8, MAPK9, AKT3, CASP3, PPP3CA, PPP3CB, PPP3R1, PPP3R2, RAF1, MAPK1, RAC1, MAPK14, MAPK11, MAPK13 | IKBKB, MAPK10, AKT2, NFKB1, RELA, FAS, TNFRSF1A, TP53, PDGFB, NFATC1, NFATC3, HRAS, MAP2K2, FOS, JUN, MYC, MAPKAPK2, MAP2K6, MAP2K7 |
| hsa04660 | T cell receptor signaling pathway | 67,78778947 | 10 | 1,31608E-17 | 1,1613E-15 | IKBKG, CHUK, AKT3, GRB2, SOS1, SOS2, PPP3CA, PPP3CB, PPP3R1, PPP3R2, CDC42, PAK1, PAK2, RAF1, MAPK1, MAP3K7, | NFKB1, RELA, IKBKB, MAP3K8, AKT2, HRAS, FOS, JUN, NFATC1, NFATC3, MAP2K2, MAP2K7 |

| | | | | | | | |
|---|---|---|---|---|---|---|---|
| | | | | | | MAPK9, MAPK14, MAPK11, MAPK13 | |
| hsa05212 | Pancreatic cancer | 71,55377778 | 10 | 1,72458E-17 | 3,04444E-15 | AKT3, RAC1, CHUK, IKBKG, BRAF, RAF1, MAPK1, MAPK8, MAPK9, CDC42, TGFB3, GADD45G | AKT2, RAC2, IKBKB, NFKB1, RELA, MAPK10, TGFB2, EGFR, TP53, GADD45A, GADD45B |
| hsa04926 | Relaxin signaling pathway | 39,05336617 | 10 | 1,8846E-17 | 1,89832E-15 | AKT3, MAPK1, ATF2, PRKACA, PRKACB, RAF1, SOS1, SOS2, MAPK14, MAPK11, MAPK13, MAPK8, MAPK9, GRB2, ARRB1 | AKT2, NFKB1, RELA, ATF4, HRAS, MAP2K2, PRKCA, FOS, JUN, MAP2K7, MAPK10, EGFR, ARRB2 |
| hsa04657 | IL-17 signaling pathway | 92,63291139 | 10 | 1,96686E-17 | 2,781E-15 | MAP3K7, CHUK, TRAF6, TAB2, IKBKG, MAPK14, MAPK1, MAPK8, MAPK11, MAPK9, MAPK13, CASP3 | IKBKB, NFKB1, RELA, FOS, JUN, JUND, MAPK10, IL1B |
| hsa05418 | Fluid shear stress and atherosclerosis | 53,6280916 | 10 | 2,18328E-17 | 8,67618E-16 | MAPK14, MAPK11, MAPK13, AKT3, MAPK8, MAPK9, RAC1, MAP3K7, CHUK, IKBKG | FOS, JUN, AKT2, MAPK10, MEF2C, MAP2K7, MAP2K6, RAC2, IL1B, PDGFB, TP53, TNFRSF1A, IL1R1, IKBKB, NFKB1, RELA, DUSP1 |
| hsa05231 | Choline metabolism in cancer | 52,93309222 | 10 | 2,88327E-17 | 3,16468E-15 | MAPK8, MAPK9, MAPK1, SOS1, SOS2, RAF1, AKT3, GRB2, RAC1 | MAPK10, HRAS, MAP2K2, AKT2, EGFR, PDGFRA, PDGFRB, PDGFB, PLA2G4A, RAC2, FOS, JUN, PRKCA, PRKCG |
| hsa05214 | Glioma | 56,37114846 | 10 | 2,88327E-17 | 3,61072E-15 | GRB2, SOS1, SOS2, BRAF, RAF1, MAPK1, AKT3, GADD45G | PDGFB, EGFR, PDGFRA, PDGFRB, PRKCA, PRKCG, MAP2K2, AKT2, TP53, HRAS, GADD45A, GADD45B |
| hsa04068 | FoxO signaling pathway | 46,0976378 | 10 | 8,75969E-17 | 1,32507E-14 | CHUK, MAPK8, MAPK9, MAPK1, SOS1, SOS2, BRAF, RAF1, AKT3, GRB2, STK4, GADD45G, NLK, MAPK14, MAPK11, MAPK13, TGFB3 | IKBKB, MAPK10, HRAS, MAP2K2, AKT2, EGFR, GADD45A, GADD45B, TGFB2 |
| hsa04915 | Estrogen signaling pathway | 42,33553719 | 10 | 1,04488E-16 | 5,54354E-15 | PRKACA, PRKACB, RAF1, MAPK1, ATF2, AKT3, GRB2, SOS1, SOS2, HSPA1L, HSPA8 | EGFR, FOS, JUN, HRAS, MAP2K2, ATF4, AKT2, HSPA2, HSPA6 |

| | | | | | | | |
|---|---|---|---|---|---|---|---|
| hsa04012 | ErbB signaling pathway | 47,3239271 | 10 | 2,09363E-16 | 2,18682E-14 | GRB2, PAK1, PAK2, MAPK8, MAPK9, SOS1, SOS2, AKT3, MAPK1, BRAF, RAF1 | ELK1, EGFR, MAP2K7, MAPK10, PRKCA, PRKCG, JUN, AKT2, MYC, MAP2K2, HRAS |
| hsa05210 | Colorectal cancer | 45,09691877 | 10 | 2,09363E-16 | 2,60333E-14 | CASP3, TGFB3, BRAF, RAF1, MAPK8, MAPK9, RAC1, MAPK1, AKT3, GADD45G, SOS1, SOS2, GRB2 | TGFB2, MAPK10, RAC2, MYC, MAP2K2, JUN, FOS, TP53, AKT2, GADD45A, GADD45B, HRAS, EGFR |
| hsa05168 | Herpes simplex infection | 46,14608513 | 10 | 2,68153E-16 | 9,03788E-13 | MAPK8, MAPK9, TAB1, TAB2, MAP3K7, IKBKG, TRAF6, CASP3, CHUK | MAPK10, NFKB1, RELA, FOS, JUN, IL1B, DAXX, TP53, FAS, TNFRSF1A, IKBKB |
| hsa04658 | Th1 and Th2 cell differentiation | 63,13568627 | 10 | 3,65032E-16 | 4,69177E-16 | CHUK, IKBKG, PPP3CA, PPP3CB, PPP3R1, PPP3R2, MAPK1, MAPK14, MAPK11, MAPK13, MAPK8, MAPK9 | NFKB1, RELA, IKBKB, NFATC1, NFATC3, MAPK10, JUN, FOS |
| hsa05166 | HTLV-I infection | 32,38604255 | 10 | 4,62002E-16 | 4,34549E-14 | PPP3CA, PPP3CB, PPP3R1, PPP3R2, PRKACA, PRKACB, ATF2, TGFB3, CHUK, IKBKG, MAPK8, MAPK9, AKT3, ELK4, MAPK1 | NFATC1, NFATC3, MYC, HRAS, ATF4, TGFB2, NFKB1, RELA, NFKB2, RELB, IKBKB, TNFRSF1A, IL1R1, MAP3K1, MAPK10, JUN, AKT2, SRF, ELK1, FOS, MAP2K2, TP53 |
| hsa05215 | Prostate cancer | 43,38735178 | 10 | 7,65225E-16 | 4,7032E-14 | CHUK, IKBKG, SOS1, SOS2, GRB2, AKT3, MAPK1, BRAF, RAF1 | NFKB1, RELA, HRAS, TP53, EGFR, FGFR2, PDGFRA, PDGFRB, IKBKB, ATF4, PDGFB, AKT2, MAP2K2 |
| hsa04664 | Fc epsilon RI signaling pathway | 55,31368103 | 10 | 7,65693E-16 | 1,0615E-13 | AKT3, MAPK14, MAPK11, MAPK13, MAPK1, RAF1, SOS1, SOS2, | AKT2, MAP2K7, MAP2K3, MAP2K6, PRKCA, MAP2K2, HRAS, PLA2G4A, MAPK10, RAC2 |

| | | | | | | | |
|---|---|---|---|---|---|---|---|
| | | | | | | GRB2, MAPK8, MAPK9, RAC1 | |
| hsa04659 | Th17 cell differentiation | 53,66533333 | 10 | 1,55151E-15 | 2,08714E-15 | CHUK, IKBKG, PPP3CA, PPP3CB, PPP3R1, PPP3R2, MAPK1, MAPK14, MAPK11, MAPK13, MAPK8, MAPK9 | NFKB1, RELA, IKBKB, NFATC1, NFATC3, MAPK10, JUN, FOS, IL1R1, IL1B |
| hsa04064 | NF-kappa B signaling pathway | 73,91919192 | 10 | 1,57854E-15 | 1,52531E-13 | CHUK, IKBKG, MAP3K7, TRAF6, TAB1, TAB2 | IL1R1, TNFRSF1A, IL1B, IKBKB, NFKB2, RELB, NFKB1, RELA, CD14, GADD45B |
| hsa04151 | PI3K-Akt signaling pathway | 18,06651176 | 10 | 2,76791E-15 | 2,77934E-13 | AKT3, NTRK2, CHUK, IKBKG, FGF9, NTF3, FGF16, GRB2, SOS1, SOS2, RAF1, MAPK1, RAC1, ATF2 | AKT2, EGFR, FGFR3, FGFR2, FGFR4, NTRK1, PDGFRA, PDGFRB, MYC, TP53, IKBKB, NFKB1, RELA, BDNF, FGF1, FGF3, FGF5, FGF7, FGF8, FGF10, PDGFB, FGF18, FGF17, FGF19, FGF20, HRAS, MAP2K2, ATF4, PRKCA, NR4A1 |
| hsa05120 | Epithelial cell signaling in Helicobacter pylori infection | 81,06092308 | 10 | 4,76011E-15 | 1,44683E-13 | MAPK14, MAPK11, MAPK13, MAPK8, MAPK9, CHUK, IKBKG, CASP3, PAK1, CDC42, RAC1 | JUN, MAPK10, IKBKB, NFKB1, RELA, EGFR |
| hsa05224 | Breast cancer | 48,14473684 | 10 | 4,90004E-15 | 7,50896E-11 | BRAF, RAF1, MAPK1, GRB2, SOS1, SOS2, AKT3, FGF9, FGF16, GADD45G | EGFR, HRAS, MAP2K2, FOS, JUN, AKT2, MYC, NFKB2, FGF1, FGF3, FGF5, FGF7, FGF8, FGF10, FGF18, FGF17, FGF19, FGF20, TP53, GADD45A, GADD45B |
| hsa04621 | NOD-like receptor signaling pathway | 49,44594595 | 10 | 6,00844E-15 | 2,32373E-11 | CHUK, MAP3K7, TAB1, TAB2, MAPK8, MAPK9, MAPK1, MAPK14, MAPK11, MAPK13, IKBKG, TRAF6 | IKBKB, IL1B, MAPK10, NFKB1, RELA, JUN |
| hsa05219 | Bladder cancer | 150,3055199 | 10 | 6,26995E-15 | 1,36367E-14 | RPS6KA5, MAPK1, BRAF, RAF1 | HRAS, EGFR, MYC, TP53, FGFR3, MAP2K2 |
| hsa04810 | Regulation of actin cytoskeleton | 45,55631013 | 10 | 6,8182E-15 | 4,27626E-13 | PAK1, PAK2, CDC42, RAC1, MAPK1, BRAF, MOS, RAF1, RRAS2, MRAS, GNA12, SOS1, SOS2, FGF9, FGF16 | RAC2, MAP2K2, HRAS, RRAS, EGFR, FGFR3, FGFR2, FGFR4, PDGFRA, PDGFRB, FGF1, FGF3, FGF5, FGF7, FGF8, |

| | | | | | | | |
|---|---|---|---|---|---|---|---|
| | | | | | | | FGF10, PDGFB, FGF18, FGF17, FGF19, FGF20 |
| hsa05218 | Melanoma | 93,37161085 | 10 | 7,52349E-15 | 6,63031E-13 | BRAF, FGF9, FGF16, AKT3, MAPK1, RAF1, GADD45G | EGFR, PDGFRA, PDGFRB, FGF1, FGF3, FGF5, FGF7, FGF8, FGF10, PDGFB, FGF18, FGF17, FGF19, FGF20, TP53, AKT2, MAP2K2, HRAS, GADD45A, GADD45B |
| hsa04622 | RIG-I-like receptor signaling pathway | 95,03896104 | 10 | 7,64468E-15 | 1,7986E-13 | TRAF6, CHUK, IKBKG, MAPK8, MAPK9, MAPK14, MAPK11, MAPK13, MAP3K7 | NFKB1, RELA, IKBKB, MAP3K1, MAPK10 |
| hsa04015 | Rap1 signaling pathway | 42,79532164 | 10 | 8,52275E-15 | 7,18846E-13 | MAPK1, RAF1, RAP1A, FGF9, FGF16, RAC1, MAPK14, MAPK11, MAPK13, BRAF, RAPGEF2, CDC42, MRAS, RASGRP2, AKT3 | MAP2K2, FGF1, FGF3, FGF5, FGF7, FGF8, FGF10, PDGFB, FGF18, FGF17, FGF19, FGF20, EGFR, FGFR3, FGFR2, FGFR4, PDGFRA, PDGFRB, RAC2, MAP2K3, MAP2K6, RASGRP3, AKT2, HRAS, RRAS, PRKCA, PRKCG |
| hsa04662 | B cell receptor signaling pathway | 90,9068323 | 10 | 1,28068E-14 | 1,20779E-11 | IKBKG, CHUK, AKT3, PPP3CA, PPP3CB, PPP3R1, PPP3R2, RAC1, MAPK1, RAF1, SOS1, SOS2, GRB2 | NFKB1, RELA, IKBKB, AKT2, FOS, JUN, HRAS, RASGRP3, NFATC1, NFATC3, RAC2, MAP2K2 |
| hsa05142 | Chagas disease (American trypanosomiasis) | 63,02583732 | 10 | 2,0441E-14 | 6,46655E-13 | TRAF6, MAPK8, MAPK9, MAPK14, MAPK11, MAPK13, MAPK1, CHUK, IKBKG, AKT3, TGFB3 | MAPK10, FOS, JUN, NFKB1, RELA, IKBKB, IL1B, TNFRSF1A, AKT2, TGFB2, FAS |
| hsa04210 | Apoptosis | 41,28102564 | 10 | 2,22467E-14 | 1,86934E-13 | CASP3, AKT3, CHUK, IKBKG, GADD45G, MAPK8, MAPK9, RAF1, MAPK1 | TP53, NFKB1, RELA, AKT2, IKBKB, NTRK1, TNFRSF1A, FAS, GADD45A, GADD45B, DAXX, MAPK10, JUN, FOS, DDIT3, ATF4, HRAS, MAP2K2 |
| hsa05226 | Gastric cancer | 56,72868217 | 10 | 2,29437E-14 | 8,05494E-11 | BRAF, RAF1, MAPK1, GRB2, SOS1, SOS2, AKT3, GADD45G, TGFB3, FGF9, FGF16 | HRAS, MAP2K2, AKT2, MYC, TP53, GADD45A, GADD45B, TGFB2, FGFR2, FGF1, FGF3, FGF5, FGF7, FGF8, FGF10, FGF18, FGF17, FGF19, FGF20, EGFR |

| | | | | | | | |
|---|---|---|---|---|---|---|---|
| hsa05223 | Non-small cell lung cancer | 45,35950413 | 10 | 3,90353E-14 | 4,35899E-12 | AKT3, STK4, SOS1, SOS2, GRB2, MAPK1, BRAF, RAF1, GADD45G | PRKCA, PRKCG, AKT2, HRAS, EGFR, MAP2K2, TP53, GADD45A, GADD45B |
| hsa04933 | AGE-RAGE signaling pathway in diabetic complications | 47,72608696 | 10 | 4,74674E-14 | 3,45795E-12 | MAPK8, MAPK9, MAPK14, MAPK11, MAPK13, MAPK1, TGFB3, CASP3, AKT3, CDC42, RAC1 | NFKB1, RELA, MAPK10, HRAS, TGFB2, IL1B, JUN, AKT2, NFATC1, PRKCA |
| hsa05220 | Chronic myeloid leukemia | 78,78229665 | 10 | 6,32224E-14 | 7,32961E-13 | TGFB3, CHUK, IKBKG, SOS1, SOS2, GRB2, AKT3, MAPK1, BRAF, RAF1, GADD45G | TP53, TGFB2, MYC, NFKB1, RELA, IKBKB, AKT2, MAP2K2, HRAS, GADD45A, GADD45B |
| hsa04917 | Prolactin signaling pathway | 45,45341615 | 10 | 6,79765E-14 | 6,39551E-12 | AKT3, MAPK1, RAF1, GRB2, SOS1, SOS2, MAPK8, MAPK9, MAPK14, MAPK11, MAPK13 | AKT2, MAP2K2, HRAS, MAPK10, FOS, NFKB1, RELA |
| hsa05131 | Shigellosis | 71,98032787 | 10 | 7,03173E-14 | 1,00107E-13 | RAC1, CDC42, IKBKG, CHUK, MAPK8, MAPK9, MAPK1, MAPK14, MAPK11, MAPK13 | IKBKB, NFKB1, RELA, MAPK10 |
| hsa04510 | Focal adhesion | 23,49277689 | 10 | 9,20877E-14 | 4,70903E-12 | CDC42, AKT3, RASGRF1, MAPK1, BRAF, RAF1, SOS1, SOS2, GRB2, RAP1A, MAPK8, MAPK9, PAK1, PAK2, RAC1 | FLNA, FLNB, FLNC, AKT2, PRKCA, PRKCG, ELK1, HRAS, EGFR, PDGFRA, PDGFRB, PDGFB, MAPK10, JUN, RAC2 |
| hsa04150 | mTOR signaling pathway | 29,56767677 | 10 | 1,19903E-13 | 2,22018E-10 | BRAF, RAF1, MAPK1, AKT3, GRB2, SOS1, SOS2, CHUK | HRAS, MAP2K2, AKT2, PRKCA, PRKCG, RPS6KA1, RPS6KA2, RPS6KA3, IKBKB, TNFRSF1A |
| hsa05164 | Influenza A | 33,75178197 | 10 | 2,32984E-13 | 3,27722E-13 | ATF2, AKT3, MAPK14, MAPK11, MAPK13, MAPK8, MAPK9, HSPA1L, HSPA8, RAF1, MAPK1 | IKBKB, NFKB1, RELA, JUN, AKT2, IL1B, MAP2K3, MAP2K6, MAP2K7, MAPK10, HSPA2, HSPA6, PRKCA, MAP2K2, FAS, TNFRSF1A |
| hsa05222 | Small cell lung cancer | 68,0392562 | 10 | 2,80696E-13 | 1,08438E-09 | AKT3, CHUK, IKBKG, TRAF6, CASP3, GADD45G | AKT2, MYC, IKBKB, NFKB1, RELA, TP53, GADD45A, GADD45B |

| | | | | | | | |
|---|---|---|---|---|---|---|---|
| hsa04062 | Chemokine signaling pathway | 21,59976387 | 10 | 3,46642E-13 | 1,01142E-09 | GRB2, ARRB1, CDC42, RASGRP2, RAP1A, PRKACA, PRKACB, PAK1, MAPK1, AKT3, BRAF, RAF1, RAC1, CHUK, IKBKG, SOS1, SOS2 | ARRB2, AKT2, HRAS, RAC2, NFKB1, RELA, IKBKB |
| hsa05133 | Pertussis | 41,50094518 | 10 | 4,62992E-13 | 1,10464E-09 | CASP3, MAPK14, MAPK11, MAPK13, MAPK1, MAPK8, MAPK9, TRAF6 | NFKB1, RELA, IL1B, FOS, JUN, MAPK10, CD14 |
| hsa04540 | Gap junction | 57,02337662 | 10 | 5,67527E-13 | 5,15737E-12 | MAP3K2, PRKACA, PRKACB, MAPK1, RAF1, SOS1, SOS2, GRB2 | PRKCA, PRKCG, MAP2K2, HRAS, EGFR, PDGFRA, PDGFRB, PDGFB |
| hsa05213 | Endometrial cancer | 94,60018467 | 10 | 5,9913E-13 | 1,08967E-11 | SOS1, SOS2, GRB2, AKT3, MAPK1, BRAF, RAF1, GADD45G | HRAS, ELK1, EGFR, MYC, AKT2, MAP2K2, TP53, GADD45A, GADD45B |
| hsa04072 | Phospholipase D signaling pathway | 57,29621575 | 10 | 1,09581E-12 | 2,0645E-12 | MAPK1, RAF1, RRAS2, MRAS, GRB2, SOS1, SOS2, AKT3, GNA12 | MAP2K2, HRAS, RRAS, AKT2, EGFR, PDGFRA, PDGFRB, PRKCA, PLA2G4A, PDGFB |
| hsa05132 | Salmonella infection | 49,40421941 | 10 | 1,10926E-12 | 5,3501E-11 | RAC1, CDC42, MAPK8, MAPK9, MAPK1, MAPK14, MAPK11, MAPK13 | NFKB1, RELA, MAPK10, FOS, JUN, CD14, FLNA, FLNB, FLNC, IL1B |
| hsa04370 | VEGF signaling pathway | 62,09212121 | 10 | 1,1794E-12 | 2,12799E-10 | CDC42, PPP3CA, PPP3CB, PPP3R1, PPP3R2, AKT3, MAPK14, MAPK11, MAPK13, MAPK1, RAF1, RAC1 | PLA2G4A, HSPB1, MAPKAPK3, MAPKAPK2, AKT2, PRKCA, PRKCG, MAP2K2, HRAS, RAC2 |
| hsa05221 | Acute myeloid leukemia | 46,77698864 | 10 | 1,98341E-12 | 8,77927E-11 | CHUK, IKBKG, SOS1, SOS2, GRB2, AKT3, MAPK1, BRAF, RAF1 | MYC, NFKB1, RELA, HRAS, IKBKB, AKT2, MAP2K2, CD14, DUSP6 |

| | | | | | | | |
|---|---|---|---|---|---|---|---|
| hsa05140 | Leishmaniasis | 77,13306983 | 10 | 1,99156E-12 | 9,51948E-10 | TGFB3, MAPK1, MAPK14, MAPK11, MAPK13, TRAF6, TAB1, TAB2, MAP3K7 | NFKB1, RELA, TGFB2, IL1B, ELK1, FOS, JUN |
| hsa04218 | Cellular senescence | 45,61080332 | 10 | 2,1125E-12 | 3,97647E-12 | AKT3, RRAS2, MRAS, RAF1, MAPK1, MAPK14, MAPK11, MAPK13, GADD45G, PPP3CA, PPP3CB, PPP3R1, PPP3R2, TGFB3 | MYC, TP53, AKT2, HRAS, RRAS, MAP2K2, MAP2K3, MAP2K6, GADD45A, GADD45B, NFKB1, RELA, MAPKAPK2, NFATC1, NFATC3, TGFB2 |
| hsa05230 | Central carbon metabolism in cancer | 49,89545455 | 10 | 3,43724E-12 | 1,98188E-11 | RAF1, MAPK1, AKT3 | MYC, EGFR, FGFR3, FGFR2, NTRK1, PDGFRA, PDGFRB, HRAS, MAP2K2, AKT2, TP53 |
| hsa04914 | Progesterone-mediated oocyte maturation | 42,64568765 | 10 | 3,49481E-12 | 5,03325E-09 | MOS, AKT3, PRKACA, PRKACB, MAPK1, MAPK14, MAPK8, MAPK11, MAPK9, MAPK13, BRAF, RAF1 | AKT2, CDC25B, RPS6KA1, RPS6KA2, RPS6KA3, MAPK10 |
| hsa04920 | Adipocytokine signaling pathway | 64,38123167 | 10 | 3,73885E-12 | 1,07001E-08 | CHUK, IKBKG, MAPK8, MAPK9, AKT3 | NFKB1, RELA, IKBKB, MAPK10, AKT2, TNFRSF1A |
| hsa04720 | Long-term potentiation | 49,89545455 | 10 | 3,87345E-12 | 1,77802E-09 | PPP3CA, PPP3CB, PPP3R1, PPP3R2, PRKACA, PRKACB, CACNA1C, MAPK1, BRAF, RAF1, RAP1A | PRKCA, PRKCG, RPS6KA1, RPS6KA2, RPS6KA3, ATF4, MAP2K2, HRAS |
| hsa04928 | Parathyroid hormone synthesis, secretion and action | 45,53422222 | 10 | 8,15972E-12 | 3,27267E-09 | PRKACA, PRKACB, ARRB1, BRAF, RAF1, MAPK1, GNA12, ATF2 | ARRB2, PRKCA, PRKCG, EGFR, ATF4, FOS, JUND, MEF2C |
| hsa04024 | cAMP signaling pathway | 43,33026316 | 10 | 1,08752E-11 | 4,32358E-10 | CACNA1C, CACNA1F, PRKACA, PRKACB, AKT3, MAPK1, RAP1A, RAF1, BRAF, MAPK8, MAPK9, RRAS2, RAC1, PAK1 | CACNA1S, AKT2, MAP2K2, MAPK10, NFKB1, RELA, NFATC1, RRAS, BDNF, FOS, JUN, RAC2 |
| hsa04071 | Sphingolipid signaling pathway | 39,02933333 | 10 | 1,16415E-11 | 5,94999E-10 | MAPK1, RAF1, AKT3, GNA12, MAPK8, MAPK9, MAPK14, MAPK11, MAPK13, RAC1 | MAP2K2, AKT2, PRKCA, PRKCG, HRAS, MAPK10, TP53, NFKB1, RELA, RAC2, TNFRSF1A |
| hsa05211 | Renal cell carcinoma | 82,95708502 | 10 | 1,3258E-11 | 1,39645E-09 | TGFB3, RAC1, SOS1, SOS2, GRB2, PAK1, PAK2, RAP1A, CDC42, AKT3, MAPK1, BRAF, RAF1 | TGFB2, PDGFB, AKT2, JUN, MAP2K2, HRAS |

| | | | | | | | |
|---|---|---|---|---|---|---|---|
| hsa04140 | Autophagy - animal | 50,10183997 | 10 | 1,67668E-11 | 2,77362E-11 | AKT3, RRAS2, MRAS, RAF1, MAPK1, MAP3K7, MAPK8, MAPK9, PRKACA, PRKACB, TRAF6 | AKT2, HRAS, RRAS, MAP2K2, MAPK10 |
| hsa05162 | Measles | 26,3997114 | 10 | 3,06655E-11 | 5,26248E-10 | AKT3, HSPA1L, HSPA8, CHUK, TRAF6, MAP3K7, TAB2 | NFKB1, RELA, AKT2, HSPA2, HSPA6, IL1B, FAS, TP53 |
| hsa04310 | Wnt signaling pathway | 35,40967742 | 10 | 4,80021E-11 | 8,29453E-11 | PPP3CA, PPP3CB, PPP3R1, PPP3R2, MAPK8, MAPK9, RAC1, NLK, MAP3K7, PRKACA, PRKACB | NFATC1, NFATC3, PRKCA, PRKCG, MAPK10, RAC2, JUN, MYC, TP53 |
| hsa05165 | Human papillomavirus infection | 14,15473888 | 10 | 5,23582E-11 | 5,43161E-09 | AKT3, CHUK, IKBKG, GRB2, SOS1, SOS2, RAF1, MAPK1, CASP3, CDC42, PRKACA, PRKACB | AKT2, EGFR, TP53, IKBKB, NFKB1, RELA, HRAS, MAP2K2, TNFRSF1A, FAS, PDGFRB |
| hsa04919 | Thyroid hormone signaling pathway | 27,46538782 | 10 | 6,66964E-11 | 1,34072E-08 | AKT3, RAF1, PRKACA, PRKACB, MAPK1 | AKT2, PRKCA, PRKCG, HRAS, MAP2K2, TP53, MYC |
| hsa04213 | Longevity regulating pathway - multiple species | 46,68580542 | 10 | 9,05017E-11 | 5,75698E-09 | AKT3, PRKACA, PRKACB, HSPA1L, HSPA8 | AKT2, HRAS, HSPA2, HSPA6 |
| hsa05225 | Hepatocellular carcinoma | 40,54293629 | 10 | 1,11294E-10 | 1,90368E-10 | BRAF, RAF1, MAPK1, GRB2, SOS1, SOS2, AKT3, GADD45G, TGFB3 | EGFR, HRAS, MAP2K2, AKT2, MYC, TP53, GADD45A, GADD45B, PRKCA, PRKCG, ELK1, TGFB2 |
| hsa05203 | Viral carcinogenesis | 33,44946673 | 10 | 1,23438E-10 | 7,37551E-09 | CASP3, IKBKG, MAPK1, ATF2, RASA2, CDC42, RAC1, PRKACA, PRKACB, GRB2 | TP53, JUN, ATF4, NFKB1, NFKB2, RELA, SRF, HRAS, MAPKAPK2 |
| hsa05216 | Thyroid cancer | 72,97150997 | 10 | 1,2489E-10 | 7,416E-09 | BRAF, MAPK1, GADD45G | HRAS, MAP2K2, MYC, NTRK1, TP53, GADD45A, GADD45B |

| | | | | | | | |
|---|---|---|---|---|---|---|---|
| hsa05152 | Tuberculosis | 27,4425 | 10 | 1,30319E-10 | 4,94845E-10 | RAF1, PPP3CA, PPP3CB, PPP3R1, PPP3R2, MAPK1, TRAF6, CASP3, AKT3, MAPK8, MAPK9, MAPK14, MAPK11, MAPK13, TGFB3 | NFKB1, RELA, CD14, TNFRSF1A, AKT2, IL1B, MAPK10, TGFB2 |
| hsa04728 | Dopaminergic synapse | 23,66582824 | 10 | 3,80282E-10 | 4,9201E-08 | PRKACA, PRKACB, CACNA1A, AKT3, ATF2, MAPK14, MAPK8, MAPK11, MAPK9, MAPK13, PPP3CA, PPP3CB, CACNA1C | PRKCA, PRKCG, CACNA1B, AKT2, ATF4, MAPK10, FOS, ARRB2 |
| hsa04910 | Insulin signaling pathway | 22,30247619 | 10 | 5,31831E-10 | 2,93999E-08 | MAPK8, MAPK9, MKNK1, MAPK1, PRKACA, PRKACB, SOS1, SOS2, GRB2, BRAF, RAF1, AKT3 | IKBKB, MAPK10, ELK1, MKNK2, HRAS, MAP2K2, AKT2 |
| hsa04650 | Natural killer cell mediated cytotoxicity | 27,27204969 | 10 | 5,81612E-10 | 1,39184E-09 | PPP3CA, PPP3CB, PPP3R1, PPP3R2, CASP3, MAPK1, BRAF, RAF1, SOS1, SOS2, GRB2, PAK1, RAC1 | NFATC1, PRKCA, PRKCG, FAS, MAP2K2, HRAS, RAC2 |
| hsa04630 | Jak-STAT signaling pathway | 24,22033097 | 10 | 8,62044E-10 | 2,75618E-07 | AKT3, SOS1, SOS2, GRB2, RAF1 | MYC, AKT2, EGFR, PDGFRA, PDGFRB, HRAS, PDGFB |
| hsa04137 | Mitophagy - animal | 47,98688525 | 10 | 1,22224E-09 | 2,15451E-08 | RRAS2, MRAS, MAPK8, MAPK9 | ATF4, RELA, TP53, HRAS, RRAS, MAPK10, JUN |
| hsa04371 | Apelin signaling pathway | 44,93508772 | 10 | 1,39645E-09 | 2,19386E-09 | AKT3, MAPK1, RRAS2, MRAS, RAF1, PRKACA, PRKACB | AKT2, MAP2K2, HRAS, RRAS, MEF2C |
| hsa04932 | Non-alcoholic fatty liver disease (NAFLD) | 36,68170426 | 10 | 3,39849E-09 | 9,6148E-08 | AKT3, CASP3, MAPK8, MAPK9, CDC42, RAC1 | AKT2, NFKB1, RELA, FAS, ATF4, DDIT3, JUN, MAPK10, IKBKB, IL1B, MAP3K11, TNFRSF1A |
| hsa04261 | Adrenergic signaling in cardiomyocytes | 21,2115942 | 10 | 5,88745E-09 | 1,28944E-07 | CACNA1C, CACNA1F, CACNA2D1, CACNB1, CACNB3, CACNG1, CACNA2D2, | CACNA1S, CACNB2, CACNG5, CACNG4, AKT2, ATF4, PRKCA |

| | | | | | | | |
|---|---|---|---|---|---|---|---|
| | | | | | | CACNG3, CACNG2, PRKACA, PRKACB, AKT3, MAPK1, ATF2, MAPK14, MAPK11, MAPK13, RPS6KA5 | |
| hsa05206 | MicroRNAs in cancer | 25,58741259 | 10 | 5,90344E-09 | 2,61741E-08 | STMN1, CASP3, MAPK1, GRB2, RPS6KA5, RAF1, SOS1, SOS2 | CDC25B, EGFR, TGFB2, MYC, HRAS, NFKB1, IKBKB, PRKCA, PRKCG, FGFR3, TP53, PDGFRA, MAP2K2, PDGFB, PDGFRB |
| hsa04723 | Retrograde endocannabinoid signaling | 21,02873563 | 10 | 6,25761E-09 | 4,30974E-05 | CACNA1A, CACNA1C, CACNA1F, PRKACA, PRKACB, MAPK14, MAPK1, MAPK8, MAPK11, MAPK9, MAPK13 | PRKCA, PRKCG, CACNA1B, CACNA1S, MAPK10 |
| hsa04115 | p53 signaling pathway | 45,03384615 | 10 | 1,05493E-08 | 0,000589801 | GADD45G, CASP3 | GADD45A, GADD45B, FAS, TP53 |
| hsa04750 | Inflammatory mediator regulation of TRP channels | 28,55406912 | 10 | 1,30006E-08 | 1,73196E-07 | PRKACA, PRKACB, MAPK14, MAPK8, MAPK11, MAPK9, MAPK13 | IL1R1, NTRK1, IL1B, PLA2G4A, MAPK10, PRKCA, PRKCG, MAP2K3, MAP2K6 |
| hsa04211 | Longevity regulating pathway | 48,67849224 | 10 | 2,23354E-08 | 1,88808E-06 | AKT3, PRKACA, PRKACB, ATF2 | AKT2, HRAS, ATF4, TP53, NFKB1, RELA |
| hsa04022 | cGMP-PKG signaling pathway | 22,00902256 | 10 | 2,26785E-08 | 0,000183051 | CACNA1C, CACNA1F, PPP3CA, PPP3CB, PPP3R1, PPP3R2, RAF1, MAPK1, ATF2, AKT3, GNA12 | CACNA1S, NFATC1, NFATC3, MEF2C, SRF, MAP2K2, ATF4, AKT2 |
| hsa04921 | Oxytocin signaling pathway | 33,86842975 | 10 | 2,84211E-08 | 5,36114E-07 | PRKACA, PRKACB, RAF1, MAPK1, CACNA1C, CACNA1F, CACNA2D1, CACNB1, CACNB3, CACNG1, CACNA2D2, CACNG3, CACNG2, PPP3CA, PPP3CB, PPP3R1, PPP3R2 | PRKCA, PRKCG, HRAS, MAP2K2, PLA2G4A, CACNA1S, CACNB2, CACNG5, CACNG4, NFATC1, NFATC3, JUN, FOS, MEF2C, ELK1, EGFR |
| hsa04066 | HIF-1 signaling pathway | 45,45341615 | 10 | 3,397E-08 | 4,22471E-05 | AKT3, MAPK1, MKNK1 | EGFR, AKT2, PRKCA, PRKCG, NFKB1, RELA, MAP2K2, MKNK2 |
| hsa04730 | Long-term depression | 72,67130089 | 10 | 5,95895E-08 | 8,25378E-08 | GNA12, CACNA1A, | PLA2G4A, PRKCA, PRKCG, MAP2K2, HRAS |

| | | | | | | | |
|---|---|---|---|---|---|---|---|
| | | | | | | MAPK1, BRAF, RAF1 | |
| hsa04931 | Insulin resistance | 35,55879495 | 10 | 5,97591E-08 | 5,18498E-06 | MAPK8, MAPK9, AKT3 | MAPK10, TNFRSF1A, IKBKB, AKT2, NFKB1, RELA, RPS6KA1, RPS6KA2, RPS6KA3 |
| hsa04934 | Cushing's syndrome | 37,27334465 | 10 | 6,05719E-08 | 1,90279E-07 | CACNA1C, CACNA1F, CACNA1H, ATF2, PRKACA, PRKACB, MAPK1, RAP1A, BRAF | CACNA1S, CACNA1I, ATF4, NR4A1, EGFR, MAP2K2 |
| hsa04520 | Adherens junction | 40,31955923 | 10 | 7,2082E-08 | 2,34946E-05 | NLK, MAP3K7, MAPK1, RAC1, CDC42 | EGFR, RAC2 |
| hsa04144 | Endocytosis | 17,66033183 | 10 | 9,6143E-08 | 1,81179E-06 | TRAF6, HSPA1L, HSPA8, CDC42, ARRB1 | HSPA2, HSPA6, HRAS, EGFR, FGFR3, FGFR2, FGFR4, PDGFRA, ARRB2 |
| hsa04623 | Cytosolic DNA-sensing pathway | 61,5993266 | 10 | 1,0614E-07 | 2,69732E-07 | CHUK, IKBKG | IL1B, NFKB1, RELA, IKBKB |
| hsa05030 | Cocaine addiction | 63,63478261 | 10 | 1,0776E-07 | 5,42014E-06 | PRKACA, PRKACB, ATF2 | JUN, NFKB1, RELA, ATF4, BDNF |
| hsa04141 | Protein processing in endoplasmic reticulum | 23,71574074 | 10 | 1,41942E-07 | 1,71563E-07 | MAPK8, MAPK9, HSPA1L, HSPA8 | ATF4, DDIT3, MAPK10, MAP2K7, HSPA2, HSPA6 |
| hsa04930 | Type II diabetes mellitus | 55,43939394 | 10 | 1,48782E-07 | 1,71563E-07 | MAPK1, MAPK8, MAPK9, CACNA1A, CACNA1C, CACNA1E | IKBKB, MAPK10, CACNA1B |
| hsa05134 | Legionellosis | 54,20740741 | 10 | 1,9485E-07 | 4,05085E-07 | CASP3, HSPA1L, HSPA8 | IL1B, NFKB1, RELA, NFKB2, HSPA2, HSPA6, CD14 |
| hsa04550 | Signaling pathways regulating pluripotency of stem cells | 34,84761905 | 10 | 2,02727E-07 | 4,06053E-06 | AKT3, GRB2, MAPK1, RAF1, MAPK14, MAPK11, MAPK13 | AKT2, HRAS, MAP2K2, FGFR3, FGFR2, FGFR4, MYC, DUSP9 |
| hsa05014 | Amyotrophic lateral sclerosis (ALS) | 48,78666667 | 10 | 2,44925E-07 | 2,794E-07 | CASP3, MAPK14, MAPK11, MAPK13, RAC1, PPP3CA, PPP3CB, PPP3R1, PPP3R2 | TNFRSF1A, DAXX, MAP2K3, MAP2K6, TP53 |
| hsa05031 | Amphetamine addiction | 32,71833085 | 10 | 2,53852E-07 | 4,89976E-07 | CACNA1C, PRKACA, PRKACB, ATF2, PPP3CA, PPP3CB, PPP3R1, PPP3R2 | ATF4, PRKCA, PRKCG, FOS, JUN |
| hsa04530 | Tight junction | 17,45151033 | 10 | 3,12113E-07 | 1,53874E-05 | CDC42, RAC1, RAP1A, PRKACA, PRKACB, MAPK8, MAPK9, RAPGEF2 | MAP2K7, MAPK10, MAP3K1, JUN |
| hsa04270 | Vascular smooth muscle contraction | 22,42492339 | 10 | 3,8048E-07 | 2,02063E-05 | GNA12, CACNA1C, CACNA1F, BRAF, RAF1, MAPK1, | PRKCA, PRKCG, CACNA1S, PLA2G4A, MAP2K2 |

| | | | | | | | |
|---|---|---|---|---|---|---|---|
| | | | | | | PRKACA, PRKACB | |
| hsa04726 | Serotonergic synapse | 54,44940476 | 10 | 4,25978E-07 | 4,72498E-06 | PRKACA, PRKACB, CACNA1A, CACNA1C, CACNA1F, CASP3, MAPK1, BRAF, RAF1 | PRKCA, PRKCG, CACNA1B, PLA2G4A, CACNA1S, DUSP1, HRAS |
| hsa05202 | Transcriptional misregulation in cancer | 21,28872727 | 10 | 5,19769E-07 | 4,66712E-05 | ELK4, GADD45G | DDIT3, MYC, CD14, MEF2C, TP53, NTRK1, DUSP6, NFKB1, RELA, GADD45A, GADD45B |
| hsa05217 | Basal cell carcinoma | 27,33146592 | 10 | 7,70421E-07 | 0,009990069 | GADD45G | TP53, GADD45A, GADD45B |
| hsa04666 | Fc gamma R-mediated phagocytosis | 19,79978355 | 10 | 1,49344E-06 | 4,19597E-05 | RAF1, MAPK1, AKT3, RAC1, CDC42, PAK1 | PLA2G4A, AKT2, RAC2, PRKCA, PRKCG |
| hsa05146 | Amoebiasis | 39,13368984 | 10 | 1,75108E-06 | 2,08636E-05 | CASP3, PRKACA, PRKACB, TGFB3 | IL1B, NFKB1, RELA, IL1R1, HSPB1, CD14, PRKCA, PRKCG, TGFB2 |
| hsa04725 | Cholinergic synapse | 20,78977273 | 10 | 2,88301E-06 | 1,57877E-05 | PRKACA, PRKACB, CACNA1A, MAPK1, AKT3, CACNA1C, CACNA1F | PRKCA, PRKCG, CACNA1B, ATF4, AKT2, CACNA1S, HRAS, FOS |
| hsa05034 | Alcoholism | 15,37394958 | 10 | 3,77301E-06 | 7,8898E-05 | PRKACA, ATF2, BRAF, RAF1, MAPK1, NTRK2, GRB2, SOS1, SOS2 | ATF4, BDNF, HRAS |
| hsa04215 | Apoptosis - multiple species | 53,80882353 | 10 | 4,95128E-06 | 8,13185E-05 | CASP3, MAPK8, MAPK9 | TNFRSF1A, MAPK10 |
| hsa04110 | Cell cycle | 21,81222057 | 10 | 5,28585E-06 | 0,000292628 | GADD45G, TGFB3 | GADD45A, GADD45B, TP53, TGFB2, CDC25B, MYC |
| hsa04670 | Leukocyte transendothelial migration | 32,00291545 | 10 | 5,38727E-06 | 0,00250711 | MAPK14, MAPK11, MAPK13, RAC1, RAP1A, CDC42 | PRKCA, PRKCG, RAC2 |
| hsa05321 | Inflammatory bowel disease (IBD) | 48,90893901 | 10 | 6,34632E-06 | 0,000531559 | TGFB3 | NFKB1, RELA, TGFB2, NFATC1, IL1B, JUN |
| hsa04611 | Platelet activation | 29,31108144 | 10 | 7,65015E-06 | 0,001122422 | RASGRP2, RAP1A, AKT3, MAPK14, MAPK11, MAPK13, MAPK1, PRKACA, PRKACB | AKT2, PLA2G4A |
| hsa04916 | Melanogenesis | 51,62610229 | 10 | 7,73856E-06 | 3,58704E-05 | PRKACA, PRKACB, MAPK1, RAF1 | PRKCA, PRKCG, MAP2K2, HRAS |
| hsa04360 | Axon guidance | 26,93411851 | 10 | 8,5554E-06 | 2,83229E-05 | MAPK1, RAC1, PPP3CA, PPP3CB, PPP3R1, PPP3R2, CDC42, PAK1, PAK2, RASA1, RAF1 | RAC2, NFATC3, HRAS, PRKCA, RRAS |
| hsa04612 | Antigen processing and presentation | 26,3997114 | 10 | 9,84458E-06 | 1,83803E-05 | HSPA1L, HSPA8 | HSPA2, HSPA6 |
| hsa04918 | Thyroid hormone synthesis | 28,18952234 | 10 | 1,03121E-05 | 0,000175567 | PRKACA, PRKACB, ATF2 | PRKCA, PRKCG, ATF4 |

| | | | | | | | |
|---|---|---|---|---|---|---|---|
| hsa04714 | Thermogenesis | 13,53752643 | 10 | 1,09955E-05 | 0,000150206 | PRKACA, PRKACB, MAPK14, MAPK11, MAPK13, ATF2, GRB2, SOS1, SOS2 | RPS6KA1, RPS6KA2, RPS6KA3, HRAS, MAP2K3 |
| hsa04911 | Insulin secretion | 24,82360923 | 10 | 1,93267E-05 | 0,000250492 | CACNA1C, CACNA1F, PRKACA, PRKACB, ATF2 | CACNA1S, PRKCA, PRKCG, ATF4 |
| hsa05032 | Morphine addiction | 19,85888738 | 10 | 2,89527E-05 | 0,000531028 | PRKACA, PRKACB, CACNA1A, ARRB1 | CACNA1B, PRKCA, PRKCG, ARRB2 |
| hsa05020 | Prion diseases | 41,57954545 | 10 | 3,00629E-05 | 0,000343665 | PRKACA, PRKACB, MAPK1 | MAP2K2, ELK1, IL1B |
| hsa04713 | Circadian entrainment | 20,78977273 | 10 | 3,32551E-05 | 7,51237E-05 | CACNA1H, MAPK1, PRKACA, PRKACB, CACNA1C, RPS6KA5 | PRKCA, PRKCG, CACNA1I, FOS |
| hsa04925 | Aldosterone synthesis and secretion | 21,88397129 | 10 | 3,32551E-05 | 7,51237E-05 | CACNA1C, CACNA1F, CACNA1H, ATF2, PRKACA, PRKACB | CACNA1S, CACNA1I, ATF4, PRKCA, PRKCG, NR4A1 |
| hsa04020 | Calcium signaling pathway | 13,85984848 | 10 | 3,7141E-05 | 0,000157879 | PPP3CA, PPP3CB, PPP3R1, PPP3R2, CACNA1C, CACNA1F, CACNA1A, CACNA1E, PRKACA, PRKACB, CACNA1H | PRKCA, PRKCG, EGFR, PDGFRA, PDGFRB, CACNA1S, CACNA1B, CACNA1I |
| hsa04961 | Endocrine and other factor-regulated calcium reabsorption | 30,94291755 | 10 | 3,87225E-05 | 0,001196663 | PRKACA, PRKACB | PRKCA, PRKCG |
| hsa04922 | Glucagon signaling pathway | 18,89979339 | 10 | 4,40505E-05 | 0,00084917 | PRKACA, PRKACB, ATF2, AKT3, PPP3CA, PPP3CB, PPP3R1, PPP3R2 | ATF4, AKT2 |
| hsa04923 | Regulation of lipolysis in adipocytes | 28,92490119 | 10 | 6,23631E-05 | 0,001660304 | PRKACA, PRKACB, AKT3 | AKT2 |
| hsa04114 | Oocyte meiosis | 16,79981635 | 10 | 0,000100989 | 0,001295953 | MOS, MAPK1, PPP3CA, PPP3CB, PPP3R1, PPP3R2, PRKACA, PRKACB | RPS6KA1, RPS6KA2, RPS6KA3 |
| hsa05416 | Viral myocarditis | 82,84528302 | 9 | 0,000124144 | 0,000124144 | CASP3, RAC1 | RAC2 |
| hsa04340 | Hedgehog signaling pathway | 33,26363636 | 10 | 0,000144093 | 0,000709646 | PRKACA, PRKACB, ARRB1 | ARRB2 |
| hsa05010 | Alzheimer's disease | 29,12636816 | 10 | 0,000167501 | 0,001723814 | CASP3, CACNA1C, CACNA1F, MAPT, MAPK1, PPP3CA, PPP3CB, PPP3R1, PPP3R2 | CACNA1S, FAS, TNFRSF1A, IL1B |

| | | | | | | | |
|---|---|---|---|---|---|---|---|
| hsa04971 | Gastric acid secretion | 24,63973064 | 10 | 0,000210537 | 0,004343474 | PRKACA, PRKACB | PRKCA, PRKCG |
| hsa04973 | Carbohydrate digestion and absorption | 48,06568144 | 10 | 0,000227657 | 0,020945681 | AKT3 | AKT2 |
| hsa04727 | GABAergic synapse | 20,15977961 | 10 | 0,000375842 | 0,006805227 | CACNA1A, PRKACA, PRKACB, CACNA1C, CACNA1F | CACNA1B, CACNA1S, PRKCA, PRKCG |
| hsa04927 | Cortisol synthesis and secretion | 38,18086957 | 10 | 0,000399138 | 0,000533083 | CACNA1C, CACNA1F, CACNA1H, ATF2, PRKACA, PRKACB | CACNA1S, CACNA1I, ATF4, NR4A1 |
| hsa04970 | Salivary secretion | 21,81222057 | 10 | 0,000406564 | 0,007195627 | PRKACA, PRKACB | PRKCA, PRKCG |
| hsa04724 | Glutamatergic synapse | 35,23916533 | 10 | 0,000413342 | 0,000822951 | CACNA1C, PRKACA, PRKACB, PPP3CA, PPP3CB, PPP3R1, PPP3R2, CACNA1A, MAPK1 | PRKCA, PRKCG, PLA2G4A |
| hsa03040 | Spliceosome | 15,36587927 | 10 | 0,00043483 | 0,000498182 | HSPA1L, HSPA8 | HSPA2, HSPA6 |
| hsa05323 | Rheumatoid arthritis | 15,52067869 | 10 | 0,000459439 | 0,008530499 | TGFB3 | IL1B, TGFB2, FOS, JUN |
| hsa04960 | Aldosterone-regulated sodium reabsorption | 40,99719888 | 10 | 0,000846385 | 0,003849108 | MAPK1 | PRKCA, PRKCG |
| hsa04217 | Necroptosis | 12,21193158 | 10 | 0,000884275 | 0,001914968 | MAPK8, MAPK9 | TNFRSF1A, FAS, IL1B, MAPK10, PLA2G4A |
| hsa05414 | Dilated cardiomyopathy (DCM) | 50,81944444 | 10 | 0,000934301 | 0,001635027 | CACNA1C, CACNA1F, CACNA2D1, CACNB1, CACNB3, CACNG1, CACNA2D2, CACNG3, CACNG2, PRKACA, PRKACB, TGFB3 | CACNA1S, CACNB2, CACNG5, CACNG4, TGFB2 |
| hsa04350 | TGF-beta signaling pathway | 24,05917808 | 10 | 0,000948298 | 0,001197449 | MAPK1, TGFB3 | MYC, TGFB2 |
| hsa05340 | Primary immunodeficiency | 107,6176471 | 3 | 0,001078726 | 0,001078726 | IKBKG | |
| hsa05412 | Arrhythmogenic right ventricular cardiomyopathy (ARVC) | 63,0862069 | 10 | 0,001108635 | 0,017382119 | CACNA1C, CACNA1F, CACNA2D1, CACNB1, CACNB3, CACNG1, CACNA2D2, CACNG3, CACNG2 | CACNA1S, CACNB2, CACNG5, CACNG4 |
| hsa05110 | Vibrio cholerae infection | 23,75974026 | 10 | 0,00115815 | 0,003141426 | PRKACA, PRKACB | PRKCA |
| hsa04260 | Cardiac muscle contraction | 62,01694915 | 10 | 0,001233964 | 0,018395294 | CACNA1C, CACNA1F, CACNA2D1, CACNB1, CACNB3, CACNG1, CACNA2D2, CACNG3, CACNG2 | CACNA1S, CACNB2, CACNG5, CACNG4 |

| | | | | | | | |
|---|---|---|---|---|---|---|---|
| hsa05410 | Hypertrophic cardiomyopathy (HCM) | 52,27142857 | 10 | 0,001321673 | 0,005625732 | CACNA1C, CACNA1F, CACNA2D1, CACNB1, CACNB3, CACNG1, CACNA2D2, CACNG3, CACNG2, TGFB3 | CACNA1S, CACNB2, CACNG5, CACNG4, TGFB2 |
| hsa04392 | Hippo signaling pathway - multiple species | 64,19298246 | 10 | 0,002195173 | 0,004301502 | PAK1 | STK3 |
| hsa04913 | Ovarian steroidogenesis | 59,73877551 | 10 | 0,00262358 | 0,004146618 | PRKACA, PRKACB | PLA2G4A |
| hsa04120 | Ubiquitin mediated proteolysis | 18,34085213 | 10 | 0,003436895 | 0,00392544 | TRAF6 | MAP3K1 |
| hsa04390 | Hippo signaling pathway | 27,1037037 | 10 | 0,003671694 | 0,004589617 | TGFB3 | STK3, TGFB2, MYC, FGF1 |
| hsa04152 | AMPK signaling pathway | 8,99017199 | 10 | 0,003923865 | 0,03459085 | AKT3, MAP3K7 | AKT2 |
| hsa05143 | African trypanosomiasis | 59,13535354 | 10 | 0,004147364 | 0,01382088 | | IL1B, FAS, PRKCA, PRKCG |
| hsa05100 | Bacterial invasion of epithelial cells | 28,25482625 | 10 | 0,004478517 | 0,019551542 | RAC1, CDC42 | |
| hsa05332 | Graft-versus-host disease | 55,75619048 | 5 | 0,005331893 | 0,005331893 | | FAS, IL1B |
| hsa04962 | Vasopressin-regulated water reabsorption | 22,17575758 | 10 | 0,005528613 | 0,014870371 | PRKACA, PRKACB | |
| hsa04940 | Type I diabetes mellitus | 51,35438596 | 5 | 0,005625265 | 0,005625265 | | FAS, IL1B |
| hsa04924 | Renin secretion | 26,08912656 | 10 | 0,00584345 | 0,013514189 | PRKACA, PRKACB, CACNA1C, CACNA1F, PPP3CA, PPP3CB, PPP3R1, PPP3R2 | CACNA1S |
| hsa04060 | Cytokine-cytokine receptor interaction | 13,742723 | 10 | 0,006418989 | 0,021598263 | TGFB3 | IL1R1, IL1B, TGFB2, FAS, TNFRSF1A |
| hsa04976 | Bile secretion | 18,10265925 | 10 | 0,010291206 | 0,032951239 | PRKACA, PRKACB | |
| hsa04972 | Pancreatic secretion | 28,64187867 | 10 | 0,010347601 | 0,027710834 | RAP1A, RAC1 | PRKCA, PRKCG |
| hsa04742 | Taste transduction | 19,71178451 | 10 | 0,012894843 | 0,042131252 | CACNA1A, CACNA1C, PRKACA, PRKACB | |
| hsa05144 | Malaria | 15,15113872 | 7 | 0,015335413 | 0,027504691 | TGFB3 | TGFB2, IL1B |
| hsa05130 | Pathogenic Escherichia coli infection | 20,84900285 | 10 | 0,017435775 | 0,021541596 | CDC42 | CD14, PRKCA |
| hsa05016 | Huntington's disease | 11,63125828 | 3 | 0,018323689 | 0,029394476 | CASP3 | BDNF, TP53 |
| hsa04145 | Phagosome | 21,52352941 | 9 | 0,022015807 | 0,022015807 | RAC1 | CD14 |
| hsa05012 | Parkinson's disease | 21,24238026 | 7 | 0,024951839 | 0,029301886 | CASP3, PRKACA, PRKACB | |
| hsa03450 | Non-homologous end-joining | 152,4583333 | 1 | 0,048090301 | 0,048090301 | | |
